# Longitudinal whole transcriptomic profiling of live cells through domain adaptation

**DOI:** 10.64898/2026.08.07.743564

**Authors:** Zhi Fei Dong, Shreya Mishra, Maha Mohamed Tageldein, Chris McIntosh, Shane M. Harding, Gregory W. Schwartz

**Affiliations:** Princess Margaret Cancer Centre, University Health Network, Toronto, ON M5G 1L7, Canada; Department of Medical Biophysics, University of Toronto, Toronto, ON M5G 1L7, Canada; Vector Institute for Artificial Intelligence, Toronto, ON M5G 1M1, Canada; Peter Munk Cardiac Centre, University Health Network, Toronto, ON M5G 1L7, Canada; Department of Computer Science, University of Toronto, Toronto, ON M5G 1L7, Canada; Department of Radiation Oncology and Immunology, University of Toronto, Toronto, ON M5T 1P5, Canada

## Abstract

Tracking transcriptomic profiles of cells over time in response to developmental cues and environmental stimuli can reveal critical insights into the fundamental mechanisms of development and disease. However, longitudinal molecular profiling at the global transcriptome level remains a major challenge, as RNA sequencing fundamentally alters or destroys cells. To overcome these limitations, we developed PENNE, a deep-learning framework that infers whole-transcriptomic profiles directly from live-cell images. Using gated attention mechanisms, PENNE trains on spatial transcriptomic datasets to align morphological features with gene expression. To enable inferences from images, our model performs domain adaptation to eliminate discrepancies between stained and unstained tissue images, effectively transferring molecular information from tissue sections to live-cell imaging. PENNE accurately identifies cell-type-specific and radiation-response markers via imputed expression. Furthermore, using only live-cell images stained with a G2/M cell cycle marker, our model captures temporal gene dynamics, evidenced by strong correlations between predicted expression and both ground-truth cellular confluency and cell-cycle progression. By bridging the gap between data-rich spatial transcriptomics and the practicality of live-cell imaging, PENNE provides a powerful new framework for monitoring molecular temporal dynamics directly through morphological information. This approach enables a paradigm-shifting workflow, fusing transcriptome-wide data with live-cell microscopy to fuel the discovery of novel gene programs via scalable, non-invasive, real-time interrogation of cellular states.

## Introduction

Cells are dynamic systems that exhibit constant changes in their molecular and functional states in response to developmental cues and environmental stimuli. A complex interplay of molecular processes, including gene expression, protein activity, and cellular signalling, creates networks and feedback loops that orchestrate the dynamics of cellular states such as the transcriptomic profiles over time. For example, during the cell cycle, cells undergo a series of tightly regulated changes in gene expression and protein activity that drive progression through different phases and checkpoints to ensure that cell division occurs in a controlled manner.^1^ This temporal dynamic of cellular states is critical for understanding development and disease, as alterations in the temporal dynamics of cellular states can lead to improper development such as cancer.^2^ However, capturing the temporal dynamics of cellular states remains a major challenge in cell biology, as the main method for profiling the molecular state of a cell, RNA sequencing, is destructive and therefore cannot be used to monitor temporal dynamics of cellular states in live cells. While technologies like MS2/PP7 RNA labelling systems^3^ are available to track the expression of specific genes in live cells, they are limited to tracking a small number of genes and may alter cellular behaviours due to the need for labelling.^4^ More recently developed methods like transcriptome storage^5^ and live-cell RNA sequencing^6^ aim to solve this issue by directly measuring the gene expression of the same cells at multiple time points, yet they are still limited by the number of time points they can track, might induce changes to existing cell mechanisms, and are currently not suitable for high-throughput applications. Therefore, these limitations create a critical gap in our ability to quantitatively track the unbiased temporal dynamics of cellular states and understand the underlying molecular mechanisms that drive these dynamics in a high-throughput manner.

While direct, non-destructive measurement of a cell’s molecular state is difficult, monitoring temporal dynamics of cellular morphology in live cells is relatively simple. Cellular morphology is another fundamental aspect of cell biology that reflects the underlying molecular and functional state of a cell. As a result, changes in cellular morphology can indicate alterations in gene expression,^7^ protein activity,^8^ and cellular responses to environmental stimuli,^9^ highlighting morphology’s critical role in understanding development and disease. For example, changes in cellular morphology in glioblastoma are associated with the transition from a neurodevelopmental to a mesenchymal state,^10^ and the degree of morphological abnormality can predict cellular survival.^11^ Therefore, monitoring the temporal dynamics of cellular morphology has become a common practice in both research and clinical settings, with applications ranging from drug screening to disease diagnosis.^12,13^ Out of the various imaging modalities available, phase contrast microscopy (PCM) is one of the most widely used techniques for monitoring cellular morphology in live cells. This popularity is due to the non-invasive properties of PCM, which enable continuous monitoring of cellular morphology without the need for staining or labelling that may alter cellular behaviour.^4^ Furthermore, compared to other non-staining methods like bright-field microscopy, PCM can produce high-contrast images due to its ability to convert phase shifts of light passing through cells into brightness changes and enhance contrast through light scattering of cell features.^14^ However, while PCM non-destructively captures morphological changes in live cells, this technique does not provide direct information about underlying molecular dynamics that drive these morphological changes. This limits the analysis of PCM images to visual inspection or simple morphological quantification such as cell size and shape, which may not fully capture the complex molecular state of the cell. Therefore, there is a critical need for methods that can bridge the gap between the morphological information captured by PCM and the underlying molecular state of the cell to enable the monitoring of the temporal dynamics of cellular states in a non-destructive and high-throughput manner.

To connect morphological information with molecular information, advancements in deep learning have enabled the creation of foundation models that can learn rich representations of cellular morphology from large-scale datasets of hematoxylin and eosin (H&E) histological images in the field of histopathology.^15–17^ The representation generated by those models can then be used for a variety of downstream tasks such as cell type classification and even direct gene expression inferences.^18–20^ The strong performance of those models have demonstrated that cellular morphology contains rich information about the underlying gene expression and that deep learning models can effectively extract this information through high-dimensional feature representations. Inspired by these works, some computational methods attempt to better quantify the celluar morphology through domain adaptation methods^21^ and even infer gene expression directly from live-cell PCM images.^22–26^ However, these methods are often limited by the number of genes they can predict or require retraining every time to adapt to new datasets. The main obstacle for such a model is a data bottleneck; while there exist rich sets of publicly-available data containing H&E-gene expression pairs through the emergence of spatial transcriptomics, there is a lack of large-scale paired PCM-gene expression datasets available for training. As a result, these methods are often trained on in-house small datasets with limited diversity using simpler machine learning models to account for smaller sample sizes, which limits their ability to generalize to new cell types and conditions. This approach of using private small datasets for training also increases the difficulty when comparing methods and evaluating performance on common benchmarks. Most importantly, such private datasets cannot be adopted by the wider community as they require users to generate their own training data and retrain their own models, which can be a major barrier for many researchers who may not have the computing resources, data, or expertise to do so.

To overcome these limitations and improve model accessibility to enable high-throughput gene expression inference of live-cell images, we introduce PENNE (Phase-to-Expression Neural Network Estimator), an open-source deep learning framework that infers whole-transcriptomic profiles directly from live-cell images. Unlike previous approaches that rely on paired PCM and gene expression datasets, PENNE leverages domain adaptation and the rich, paired H&E-gene expression datasets from spatial transcriptomics to transfer molecular knowledge learned from H&E images to live-cell PCM images. This process enables us to apply PENNE to large and diverse datasets and generalize to new cell types and conditions without the need for retraining. We show PENNE’s ability to accurately infer cell type, radiation damage response, confluency, and cell cycle-specific gene expression markers from four independent PCM image datasets with different cell types and conditions, demonstrating PENNE’s broad applicability to find novel gene expression dynamics directly from live-cell PCM images. The PENNE framework and a pre-trained model are freely available at https://github.com/schwartzlab-methods/penne.

## Results

### PENNE infers live-cell gene expression by adapting H&E images from spatial transcriptomic assays

PENNE is a deep learning framework that infers whole transcriptomic profiles directly from live-cell PCM images, enabling new workflows such as real-time identification of cell types as well as monitoring cell states such as confluency, cell cycle, and radiation-induced damage (Figure 1a). We trained PENNE using paired gene expression and cell morphology data from spatial transcriptomic H&E images from public 10x Genomics Visium datasets with 46,161 paired H&E patches and spatial transcriptomic data across nine different samples,^27^ where the model learned to map morphological features to gene expression profiles (Figure 1b). To enable inference on PCM images, PENNE uses features extracted from large-scale H&E foundation models by converting PCM images to H&E-like images with SPAGHETTI.^21^ PENNE then performs domain disentanglement to separate the domain-invariant “biological” features from the “domain”-specific features and performs domain adaptation to further align the domain-invariant biological features between H&E and PCM images beyond SPAGHETTI’s conversion. PENNE then refines the gene-morphology mapping with constraints from known marker genes on cell lines from the LIVECell^28^ dataset. The trained model effectively transfers the molecular knowledge learned from spatial transcriptomic H&E images to PCM images, enabling high-throughput and accurate gene-expression inference directly from live-cell imaging.

**Figure 1:**
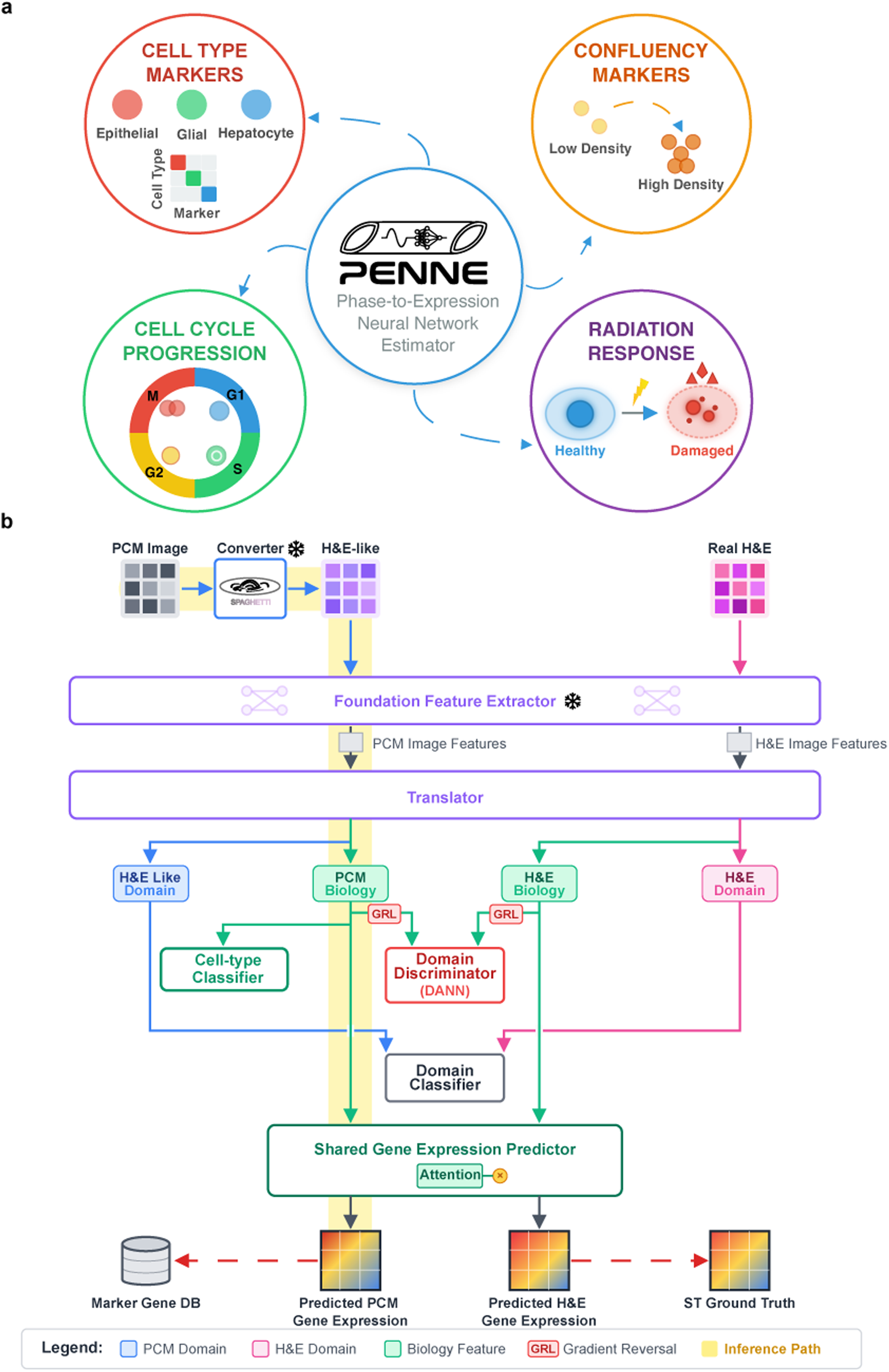
Overview of Phase-to-Expression Neural Network Estimator (PENNE). **a**, Demonstrated use cases of PENNE. PENNE is a deep learning model that predicts gene expression directly from phase contrast microscopy (PCM) images to identify cell types and track confluency, radiation, and cell-cycle-related markers, enabling new molecular insights directly from PCM. **b**, Overview of the PENNE architecture. PENNE leverages a domain adaptation framework to learn a shared latent space between PCM live-cell morphologies and H&E tissue morphologies, enabling the model to learn the mapping from H&E features to gene expression profiles and then adapt this mapping to PCM images. PENNE consists of pretrained blocks that extract morphological features from images,^16,21^ and trainable blocks that perform domain adaptation and gene-expression inference using by splitting “domain”-associated features out from “biology” features, which feed into a regression to infer gene expression using spatial transcriptomic datasets.

### PENNE accurately infers live-cell type-specific gene expression markers

To test the accuracy of PENNE’s gene expression inference on PCM images, we first evaluated whether PENNE could accurately infer the expression of cell type-specific marker genes from PCM images. Currently, identifying cell types from PCM images relies on manual annotation by experts or labelling via fluorescent markers.^29^ However, manual annotation is time-consuming and may be subject to inter-observer variability, and fluorescent labelling is labour-intensive and may alter cell behaviour.^4^ While there also exist computational methods for cell-type classification,^30^ their classification abilities are limited to the cell types within training sets and cannot be used on other cell types that the model is not trained on. PENNE presents an orthogonal approach to live-cell-type identification by inferring gene expression profiles from PCM images and determining cell type based on the expression of known marker genes. After applying PENNE to PCM images of six different human cell lines (Huh7 hepatocellular carcinoma, A172 glioblastoma, BT474 breast cancer, MCF7 breast cancer, SkBr3 breast cancer, and SkOV3 ovarian cancer cell lines) from the testing portion of the LIVECell dataset,^28^ we found that PENNE accurately inferred the expression of cell-type-specific marker genes indicated by significantly higher expression of the marker genes in the correct cell line compared to other cell lines (two-tailed Mann-Whitney U test: all *p* < 2.26 × 10^−16^; Figures 2a and 2b). As deep learning models can sometimes rely on shortcut learning of hidden data acquisition biases and result in over-promising benchmark performances,^31^ we investigated whether PENNE relies on the expected cellular morphologies in the PCM images to make accurate gene-expression inferences. We showed that morphological features are indeed important for PENNE, as using randomly permuted PCM images with destroyed morphological features as input would lead to a worse cell-type classification performance when comparing the Area Under the Receiver Operating Characteristic (AUROC) curves by using the predicted expression of marker genes to classify the cell types (Supplementary Figure S1). Surprisingly, we found that PENNE accurately inferred the expression of cell-type-specific marker genes for images of the U373 glioblastoma cell line,^32^ even though U373 was not included in training (two-tailed Mann-Whitney U test: *p* < 2.26 × 10^−16^; Figure 2c). To determine if these results were found by chance, we randomly permuted the input PCM images. With this corrupted dataset, PENNE could no longer infer the expression of cell type-specific marker genes (two-tailed Mann-Whitney U test: *p* < 2.26 × 10^−16^; Figure 2c). Taken together, these results demonstrate that PENNE relies on morphological features in the PCM images to make accurate gene-expression inferences for one-shot transference to new cell types and datasets not seen during training.

**Figure 2:**
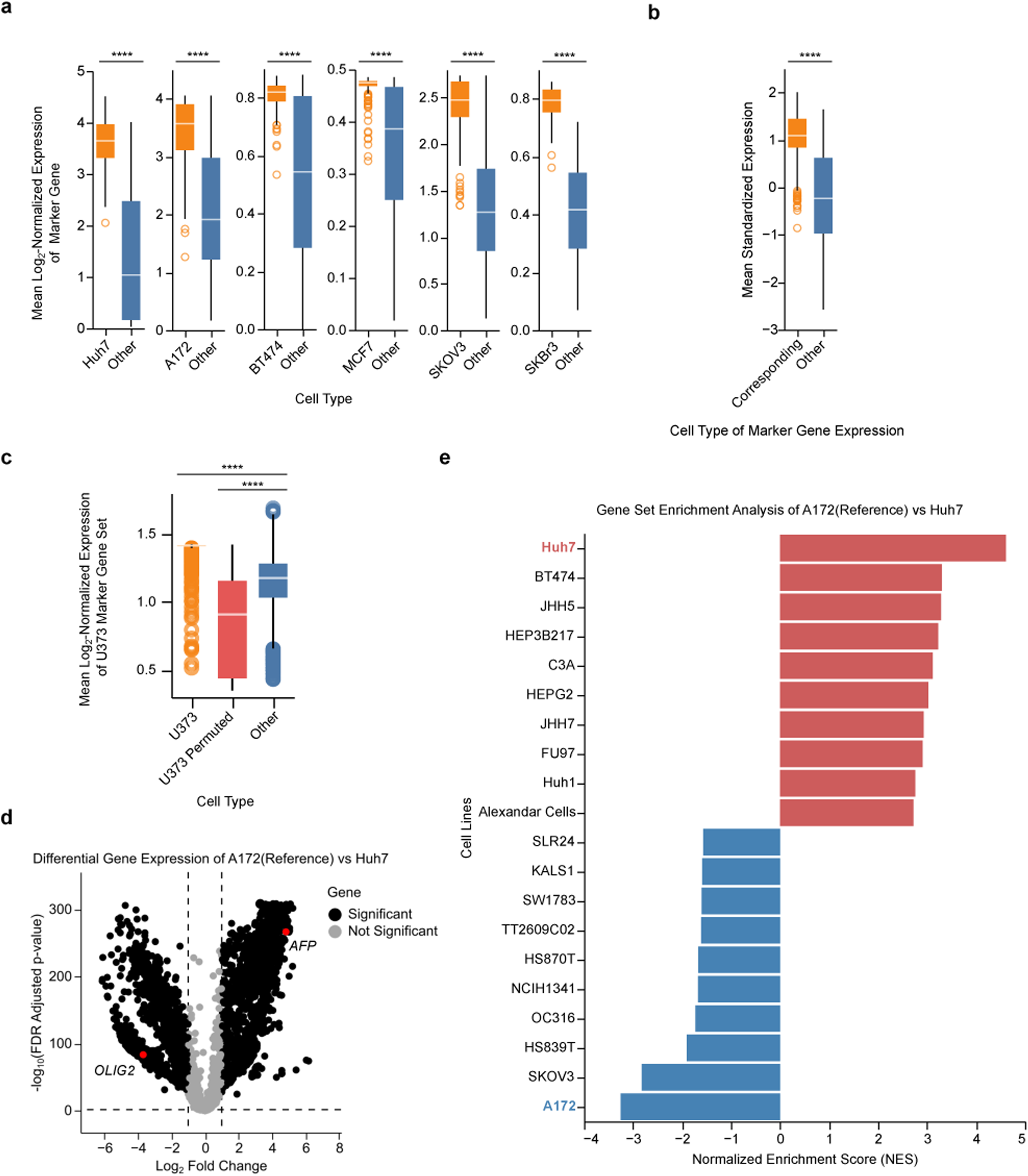
PENNE predicts cell-type-specific marker gene expression from live-cell images. **a**, Box-and-whisker plots (center line, median; box limits, upper (75^th^) and lower (25^th^) percentiles; whiskers, 1.5 × interquartile range; points, outliers) showing the mean log_2_ normalized gene expression levels of marker genes for each cell type compared to all other cell types in the LIVECell dataset.^28^ In all cases, the marker gene expression for the respective cell types are significantly higher than for other cell types. **b**, Box-and-whisker plot of the overall mean standardized expression levels of the marker gene expression profiles for all cell types in their corresponding cell type compared to those for all cell types in other cell types. The corresponding marker expression is significantly higher than expression in other cells, suggesting that PENNE captures biologically-relevant expression based on cell-type morphology. **c**, Box-and-whisker plot of the mean log-counts-per-million normalized expression levels of the marker gene expression profiles for U373 cells in the original U373 cell images^32^ compared to those for randomly permuted U373 cell images and all cell types in the LIVECell PCM images. Despite the omission of U373 cells from the training data, the marker gene expression profiles for U373 cells are significantly enriched in U373 cell images. **d**, Volcano plot showing the differentially expressed genes between A172 cell images and Huh7 cell images. Key glioblastoma marker genes such as *OLIG2* are significantly upregulated in A172 cell images, while key hepatocellular carcinoma marker genes such as *AFP* are significantly upregulated in Huh7 cell images. **e**, GSEA plot showing the significant enrichment of different cell-line-related gene sets^91^ in the differentially-expressed genes between A172 cell images and Huh7 cell images. Both Huh7 and A172 cell images show significant enrichment of gene sets related to their respective cell lineage (highlighted in blue and red, respectively). Two-tailed Mann-Whitney U test, ****: *p* < 1 × 10^−4^.

We next inquired whether PENNE expression could detect transcriptome-wide, cell-type-specific gene expression and gene-expression programs. A differential gene expression (DEG) analysis on all combinations of the six cell lines revealed that predicted gene-expression profiles showed key marker genes significantly upregulated in the corresponding cell types. For example, when comparing the A172 to Huh7 cell lines, we found that the glioblastoma marker genes (e.g. *OLIG2*^33^) were significantly upregulated in A172 PCM images compared to Huh7 PCM images, and hepatocellular carcinoma marker genes (e.g. *AFP*^34^) were significantly upregulated in Huh7 PCM images compared to A172 PCM images (Student’s *t*-test, adjusted *p* < 0.05, log2 fold change > 1; Figure 2d). Similarly, gene set enrichment analysis (GSEA)^35^ of the inferred expression revealed that both A172 and Huh7 were enriched for their corresponding gene sets (Figure 2e). This strong enrichment of cell-type-specific gene expression programs in the predicted gene expression profiles was consistent across 27 out of 30 combinations of the six cell lines (Supplementary Figure S2), suggesting that PENNE can distinguish between cell types with similar origins such as MCF7 and SkBr3 breast cancer cell lines.

We further validated PENNE’s ability to identify cell type-specific gene expressions by creating a 1:1 mixture of MCF10A normal breast epithelial cells transduced with H2B-GFP and HCT116 colorectal cancer cells, neither of which were included in the LIVECell training dataset. We generated the gene-expression profiles from the PCM images using PENNE and measured the green fluorescence intensity of the PCM images to represent number of MCF10A cells in the image (Figure 3a). We found that the predicted gene expression accurately predicted the MCF10A fraction with a significantly strong positive correlation through a Ridge regression, suggesting that the inferred gene-expression profiles relate to the presence or absence of specific cell types (Coefficient of determination: *R*^2^ = 0.510, *F* test: *p* < 2.26 × 10^−16^; Figure 3b). When investigating the genes most positively-associated with the regression, we found significant enrichment for cellular stability pathways such as the p53 pathway and apical junction found in non-tumoural epithelial MCF10A cells^36^ (Fisher’s exact test, adjusted *p* < 0.250; Supplementary Figure S3). In contrast, the most negatively-associated genes are significantly enriched for cellular stress and growth pathways such as hypoxia and mitotic spindle found in tumoural HCT116 cells^37^ (Fisher’s exact test, adjusted *p* < 0.250).

**Figure 3:**
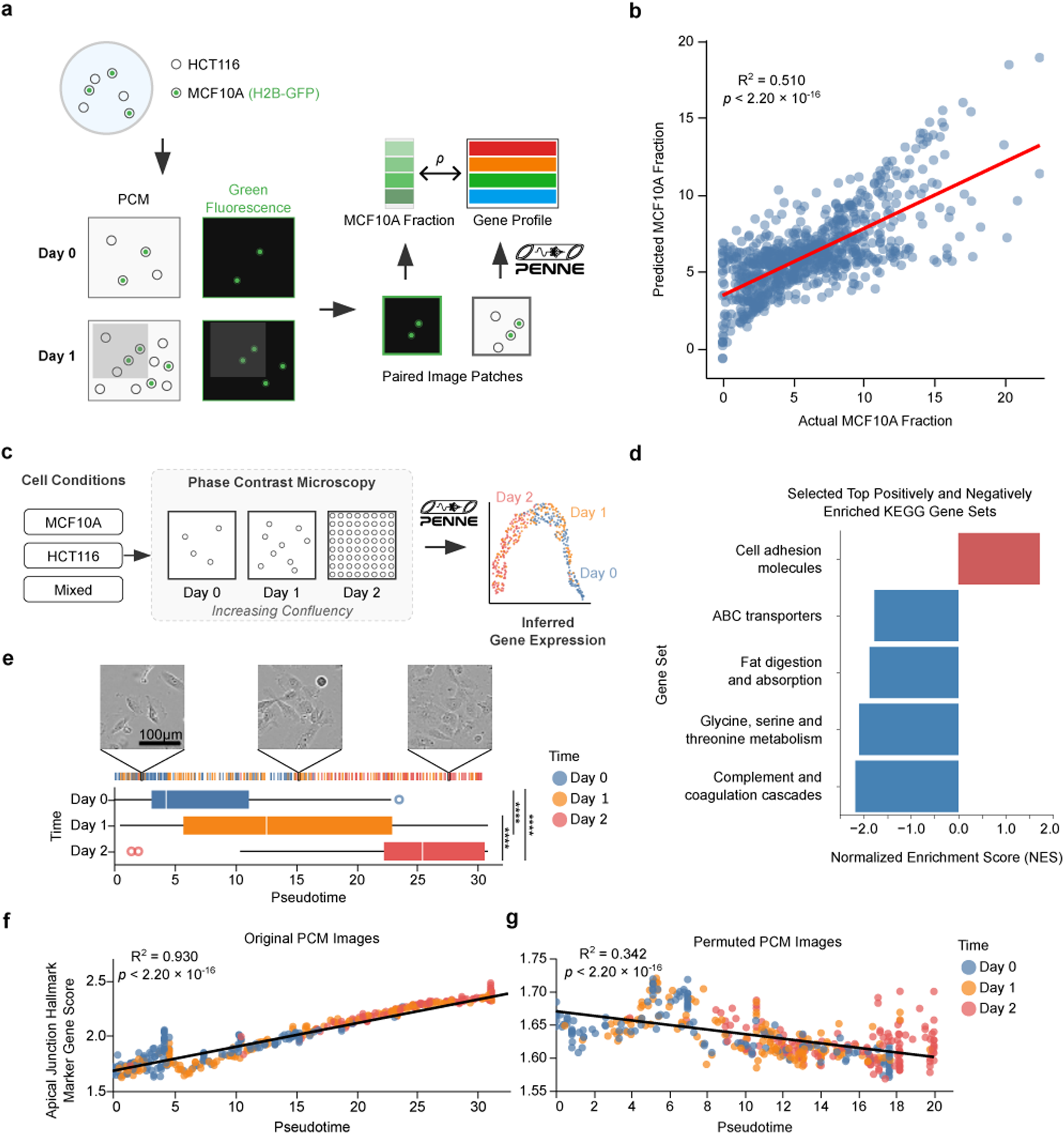
PENNE successfully predicts cell-type-specific marker gene expression from co-culture and confluency-related gene-expression profiles from live-cell PCM images. **a**, Workflow of the MCF10A (H2B-GFP) and HCT116 co-culture experiment. We used green fluorescence as a measure of MCF10A-cell abundance in each image. We then generated gene expression from PENNE for each PCM image and built a ridge regression model to predict the green fluorescence intensity from the predicted gene expression profiles. **b**, Scatter plot of the predicted MCF10A cell fraction from a Ridge regression model using PENNE-predicted gene-expression profiles compared to the actual fractions of MCF10A cells in the images reported by green light intensity. **c**, Workflow of the confluency experiment. We generated gene-expression profiles using PENNE on PCM images across three time points and compared the differences in gene expression between different confluencies. **d**, GSEA analysis showing significant enrichment of KEGG pathways^102^ in the differentially-expressed genes between day 2 (high confluency) and day 0 (low confluency). **e**, Examples and tick plots showing pseudo-time ordering of the PCM images (top) and the distribution of images from different days along pseudo-time (bottom). We observed a significantly higher pseudo-time for images later in the time course, where cells are more confluent. The example PCM images are enhanced for better visualization of the cell morphologies and confluencies. **f**, **g**, Scatter plots showing the mean expression levels of the MSigDB Hallmark^100^ Apical Junction gene set in the original PCM images (**f**) and the permuted PCM images (**g**) compared to pseudo-time. We observed a significant positive correlation in the original images that is disrupted with a significantly negative correlation in the permuted PCM images. Two-tailed Mann-Whitney *U* test, ****: *p* < 1 × 10^−4^.

To determine whether cell types from our PENNE-inferred gene expression correlated with ground-truth RNA-seq, we compared the difference between MCF10A and HCT116 marker genes from either the predicted expression or from bulk-RNA sequencing data for these cell lines. We found that patches with a lower fraction of MCF10A (Supplementary Figure S4a) had a lower difference of expression compared to patches with a higher MCF10A fraction (two-tailed Mann-Whitney U test: *p* = 0.0764; Supplementary Figures S4b and S5a). When correlating different levels of MCF10A fraction with the difference of marker gene expressions, we also found that there is a significant positive correlation between the two, suggesting that PENNE is predicting MCF10A and HCT116 marker gene expression in a way that correlates with the ground population of the two cell types in the images (coefficient of determination: *R*^2^ = 0.442, *F* test: *p* = 0.0358; Supplementary Figure S5b). Together, these results suggest that PENNE accurately infers cell-type-specific gene expression from PCM images, both in individual cell lines and in mixtures of cell lines not previously seen by PENNE. These results position the model as a powerful one-shot method for cell-type identification directly from live-cell imaging without the need for manual annotation or fluorescent labelling.

### PENNE identifies confluency-dependent gene-expression changes in longitudinal samples

After validating cell type classification capabilities using PENNE-inferred gene expression, we tested whether PENNE could report the temporal dynamics of cellular states such as cellular confluency and radiation. Confluency affects multiple critical cellular behaviours including proliferation.^38^ Yet, as sequencing is destructive, current approaches that aim to understand transcriptional differences across confluencies are limited to comparing different sample snapshots.^39^ These snapshots may introduce confounding factors such as batch effects. Here, we sought to use PENNE’s ability to infer real-time confluency-dependent gene expression changes directly from live-cell PCM images. We used PENNE to generate gene-expression profiles of MCF10A and HCT116 cells at day 0 and day 2 after seeding when the cells are at low confluency and high confluency, respectively (Figure 3c). Pathway enrichment between day 0 and day 2 revealed upregulation of cell adhesion pathways at day 2, while cells at day 0 were significantly enriched for gene sets related to metabolism, consistent with the expected biology of cells at different confluencies^40,41^ (Fisher’s exact test, adjusted *p* < 0.250; Figure 3d). In addition, cells at day 2 were significantly downregulated for gene sets related to complement and ABC transporters (Fisher’s exact test, adjusted *p* < 0.250), perhaps due to the behaviour of cells immediately after seeding, where the cellular stress caused by the new environment^42^ and extra-cellular matrix detachment through proteases drives up expressions of drug-resistance pumps like ABC transporters^43,44^ and complement components.^45^

To further investigate the temporal dynamics of gene-expression changes associated with cellular confluency, we projected the gene expression profiles to a Uniform Manifold Approximation and Projection (UMAP) embedding and found that day 0 and day 2 images were well separated with day 1 images in the middle (Supplementary Figure S6), suggesting a valid trajectory from predicted gene-expression profiles. To validate this finding, we conducted a pseudo-time analysis using the predicted gene-expression profiles from PCM images to order the images along a trajectory of cellular confluency. We found that day 0 and day 2 images were well separated along the pseudo-time trajectory with day 1 images in between, and we observed significantly higher average pseudo-times as the cellular confluency increases from day 0 to day 2, as expected (two-tailed Mann-Whitney *U* test: between day 0 and day 1 *p* = 9.15 × 10^−12^, all other comparisons *p* < 2.26 × 10^−16^; Figure 3e). For orthogonal validation, we measured the mean expression of a key confluency marker gene set, the apical junction,^46^ across cellular confluency and pseudo-time. We found a strong positive correlation between the mean expression of apical junction genes and the pseudo-time (coefficient of determination *R*^2^ = 0.930, *F* test: *p* < 2.26 × 10^−16^; Figure 3f) along with a significant correlation between mean expression value and cellular confluency (two-tailed Wilcoxon rank-sum test: between day 0 and day 1 *p* = 1.20 × 10^−5^, all other comparisons *p* < 2.26 × 10^−16^; Supplementary Figure S7a). This relationship becomes strikingly weaker (coefficient of determination *R*^2^ = 0.342, *F* test: *p* < 2.26 × 10^−16^; Figure 3g) as expected, and the mean expression decreases as the cells become more confluent (Supplementary Figure S7b) for permuted PCM images, suggesting that PENNE tracks the true temporal dynamics of cellular confluency through the underlying morphological features in the PCM images. We also found that the mean expression of apical junction genes is significantly higher in MCF10A cells compared to HCT116 cells (two-tailed Wilcoxon rank-sum test: *p* = 2.45 × 10^−9^) and mixed MCF10A/HCT116 cells (two-tailed Wilcoxon rank-sum test: *p* = 0.0130; Supplementary Figure S8) at day 2. This is consistent with the fact that MCF10A cells are normal epithelial cells and therefore have a stronger cell-cell adhesion compared to HCT116 cancer cells,^47,48^ further confirming that PENNE can capture the temporal dynamics of cellular confluency by inferring the expression of key confluency marker genes from live-cell PCM images.

### PENNE identifies radiation related gene-expression changes in perturbation experiments

Radiation therapy is one of the most common methods for treating solid tumours and is used in about 50% of all cancer patients during the course of the illness.^49,50^ Existing studies have shown that various factors that change over time such as cell cycle phases^51^ and stages of DNA damage repair^52^ can affect the radiation response of cells, and therefore improving treatment strategies requires the understanding of gene expression changes temporally. However, while past studies have demonstrated the relationship between radiation response and temporal dose dynamics,^53,54^ such findings were limited by the destructive nature of sequencing and therefore cannot capture the temporal gene-expression dynamics of radiation response of the same cellular population, potentially leading to survivorship biases where only surviving cells at the end of the experiment are analyzed. To overcome this limitation and to evaluate whether PENNE can capture the temporal dynamics of radiation response as opposed to previously unperturbed confluence, we then applied PENNE to temporal PCM images of HCT116 and MCF10A cells at day 0 and day 2 after 10 Gy of ionizing radiation (Figure 4a). We used TooManyCells^55^ divisive hierarchical clustering and visualization with TooManyCells Interactive^56^ on PENNE-predicted gene-expression profiles and identified distinct clusters of irradiated and non-irradiated patches as well as MCF10A- and HCT116-dominant patches (Figure 4b). Interestingly, we observed that cell types better separated in non-irradiated clusters compared to the irradiated clusters, suggesting that radiation may induce a more similar gene expression program in both cell types when they activate damage-response pathways. Differential gene expression followed by pathway enrichment between the irradiated groups and non-irradiated groups for both MCF10A and HCT116 cells reported that while both cell types exhibited similar damage response pathways such as UV response and complement, only MCF10A cells exhibited significant up regulation of p53 pathway (Fisher’s exact test, adjusted *p* < 0.250; Figure 4c). In contrast, HCT116 cells relied more heavily on other damage response pathways such as TGF-beta signalling (Fisher’s exact test, adjusted *p* < 0.250; Figure 4d). As the imaged HCT116 cells were p53 null and therefore may rely on other p53 independent pathways to respond to radiation-induced damage, we determined that PENNE may be identifying p53-associated morphological features that translated into associated gene expression.^57^ Overall, these results demonstrate that PENNE-powered analyses can monitor the temporal dynamics of radiation response directly from live-cell PCM images, providing a powerful method for understanding the molecular mechanisms underlying radiation response in order to optimize temporal treatment strategies.

**Figure 4:**
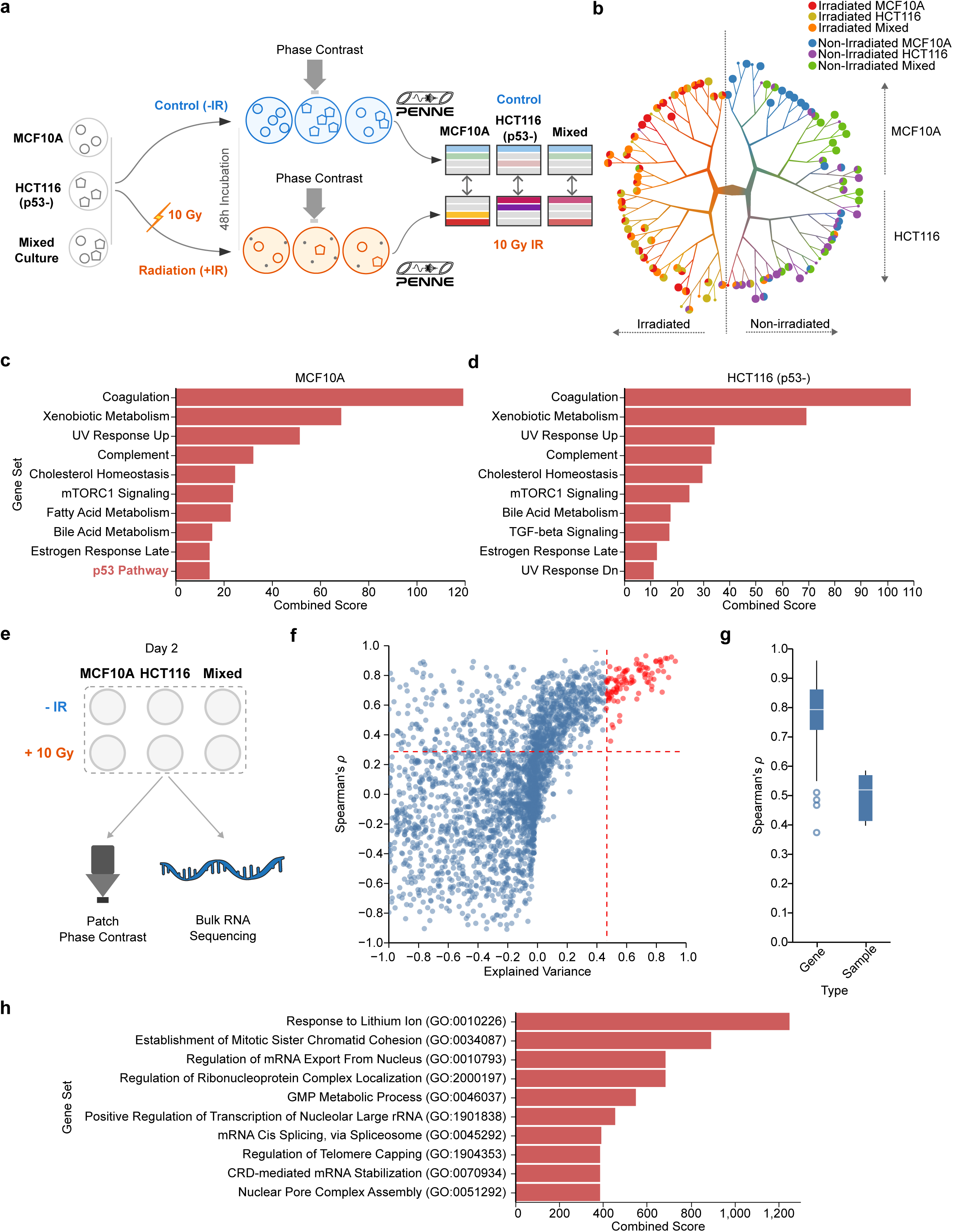
PENNE infers radiation-response-related gene expression from PCM images and identifies high-confidence genes associated with cellular structure. **a**, Workflow of the radiation response experiment. We compared the inferred gene-expression profiles from PCM images of MCF10A and HCT116 p53-null cells for non-irradiated (NIR) and 10 Gy irradiated (IR) conditions to identify radiation-response-related gene-expression profiles. **b**, TooManyCells^55^ clustering dendrogram of gene-expression profiles inferred from PCM images. Images generally clustered by treatment condition and cell type. **c**, **d**, Enrichment analysis using the MSigDB Hallmark gene sets^100^ of the most positively-correlated genes from MCF10A (**c**) and HCT116 p53-null (**d**) cell lines. While both cell lines have significant enrichment for radiation-damage-related pathways, only MCF10A has significant enrichment for the p53 pathway (red highlight).**e**, Workflow identifying high-confidence genes using ground-truth bulk RNA-sequencing data. We compared the pseudo-bulk-predicted gene-expression profiles from PCM and the ground-truth bulk RNA-sequencing data across all samples to identify high-confidence genes. **f**, Scatter plot showing the mean Spearman’s *ρ* and explained variance between the pseudo-bulk-predicted gene-expression profiles from PCM images and the bulk RNA-sequencing data for each gene across all samples. We defined “high confidence” genes (red) as genes with a mean Spearman’s *ρ* greater than 0.3 and an explained variance greater than 0.5 (dashed lines: thresholds). **g**, Box-and-whisker plot showing the gene and sample correlation between the ground-truth and predicted gene-expression profiles using high confidence genes. **h**, Enrichment analysis using Gene Ontology gene sets^103^ of the high confidence genes. The enriched major pathways relate to cell cycle and cell division.

### Reliable PENNE gene-expression predictions use morphological features of different cellular organelles

After validating PENNE’s gene-expression inference capability, we hypothesized distinct sets of genes would benefit differently from cellular morphology as an input into the model. As such, we identified a set of high-confidence genes by comparing the predicted expression profiles from PCM images to ground-truth gene-expression profiles from bulk RNA sequencing data across all 18 samples (Figure 4e). Through this process, we determined 92 genes (Supplementary Note S1) to be high confidence genes with not only a high correlation but also a high explained variance between the predicted and ground-truth gene-expression profiles across all samples (Figure 4f). Furthermore, we found that this set of genes is also highly correlated within each sample (Figure 4g). As a result, this high-confidence gene set is reliable for downstream analyses, potentially avoiding deep-learning model pitfalls such as mean and mode collapse — where a model may predict similar gene-expression profiles across all samples when training on high-dimensional and sparse data.^58^

Enrichment analysis on this set of genes reported pathways related to cell division and cell-cycle progression (Figure 4h). This finding is consistent with other reports stating that cell-cycle-related programs are better reflected in cellular morphology in imaging, promoting accurate inferences.^59^ Gating attention correlation pattern of PENNE’s Gene Expression Predictor Module also suggested that our model groups genes into clusters related to different cellular organelles such as nucleus, bounding membrane, and trans-Golgi network that are visible from PCM images^60^ (Supplementary Figures S9a and S9b). This enrichment completely disappears on random permutation of PCM images (Supplementary Figure S10), which may suggest that the gating attention module relies on biologically meaningful morphological information from organelles to assist PENNE’s accurate gene-expression inferences for genes related to cell division and cell cycle progression.

### Domain adaptation, biological constraints, and specialized losses most contribute to gene-expression inference

To evaluate the importance of each component of PENNE for gene-expression inference from live-cell PCM images, we performed an ablation experiment including adversarial domain adaptation, biological constraints, and specialized non-zero constraint losses (Supplementary Table S1). We found that while removing the adversarial domain adaptation component achieves better performance when compared to using the complete PENNE model on all genes, the complete PENNE model still achieves a higher correlation compared to all other ablations when in the context of genes with non-zero expression in ground-truth data and using our high-confidence gene set. We also observed that the performance of the model in all ablation cases drops while the performance of the complete version of PENNE increases when we evaluate on only non-zero genes and high-confidence genes. Furthermore, we found that the experiment where eliminating the non-zero constraint loss has the largest impact on the performance of the model. This finding suggests that the non-zero constraint loss is critical for preventing mode collapse as removing this loss would lead to a significant drop in the number of genes with non-zero predicted expression (two-tailed Mann-Whitney *U* test, *p* < 2.26 × 10^−16^; Supplementary Figure S11). Therefore, all of PENNE’s components are critical for accurate gene expression inference from PCM images.

### PENNE predicts periodic cell-cycle-related gene expression directly from live-cell images and identifies novel cell-cycle-related gene-expression dynamics

Given that many of our model’s high-confidence genes are involved in the cell cycle and mitosis, we used PENNE to track cell-cycle progression directly from live-cell PCM images. Cell-cycle progression is driven by a complex interplay of molecular and cellular processes, heavily orchestrated by changes in gene expression. Tracking cell-cycle progression and dysregulation is critical for understanding development and disease. For example, cancer is often characterized by uncontrolled cell division, and many anti-cancer drugs target specific phases of the cell cycle to inhibit cancer growth.^61^ However, current methods for tracking cell cycle progression often rely on fluorescent markers, which limit the number of genes that can be tracked simultaneously and may alter cell behaviour.^4^ One of these fluorescent markers is geminin, a DNA replication inhibitor whose expression level is closely related to cell cycle progression: geminin expression is low during G1 phase, increases through S phase, and peaks at G2/M phase to promote cell division.^62^ While useful to track cell-cycle dynamics, such many such markers are difficult to simultaneously track and prohibit discovery of new biology. These constraints create obstacles in measuring unknown cell-cycle-related gene-expression dynamics. With PENNE, we sought to bypass these impediments by inferring the gene expression of key cell-cycle genes, and comparing them to ground-truth geminin fluorescence intensity.

To determine whether PENNE-predicted gene-expression levels can track the temporal dynamics of cell-cycle progression, we conducted a cross-correlation analysis between PENNE output levels of each high-confidence gene with geminin fluorescence intensity across different time lags (Figure 5a). The cross-correlation heatmap revealed five distinct clusters of genes with periodic patterns, reflecting a cyclic behaviour (Figure 5b and Supplementary Figure S12). Using fast Fourier transform,^63^ we found the mean intensity frequency of geminin signal was mainly ≈0.05, or a periodicity of ≈20 h (Figure 5c), consistent with the expected overall cell cycle length of MCF10A.^64^ When we repeated the analysis on MCF10A cells that were irradiated with 5Gy of radiation, we found that the dominant frequency of the geminin signal decreases to a frequency of around 0.020 (Supplementary Figure S13), consistent with the expected lengthening of cell cycle progression after radiation treatment.^51^ Yet in both conditions, most clusters have a similar frequency to that of geminin, suggesting that the predicted gene expression levels of those genes are closely tracking the temporal dynamics of cell cycle progression. To further investigate the biological relevance of these clusters from the non-treated group, we examined the transcription factor targets enriched in each cluster. We found that the genes in clusters 1 and 2 also had a peak periodicity at ∼20 h and are significantly enriched for transcription factors IRF9 and CTCF, respectively (Figure 5d). IRF9 is a key transcription factor mediating the interferon response plays a critical role in inducing anti-proliferative and pro-apoptotic gene-expression programs in response to cellular stress in the G1/S phase.^65,66^ In contrast, CTCF is involved in regulating chromatin architecture during the G2/M phase of the cell cycle.^67^ This inverse relationship aligns with the correlation pattern of clusters 1 and 2: cluster 1 genes negatively correlate with geminin, involving genes inhibiting cell-cycle progression (*CDKN1A* and *CDKN1B*^68^) during a time of low geminin level, while the cluster 2 genes positively correlate and are involved in chromatin and mitotic spindle regulation to promote cell-cycle progression (*SPDL1* and *HNRNPU*^69,70^). Interestingly, cluster 3 had a lower periodicity at ≈10 h and is significantly enriched for transcription factor NKX2-1. Similar to geminin, NKX2-1 is a key regulator of proliferation and cell cycle progression. Yet unlike geminin, the targets of NKX2-1, including *SP1* that regulates the expression of many cell cycle regulators such as Cyclin D1,^71,72^ are elevated in expression in both G1 where geminin levels are low and G2 where geminin levels are high.^73^ Therefore, the predicted gene expression levels of Cluster 3 are positively correlated with geminin at lag 0 as those genes are involved in promoting cell cycle progression during a time of high geminin level in G2, while also having a lower periodicity at around 10 hours due to their other elevated expression peak during G1.

**Figure 5:**
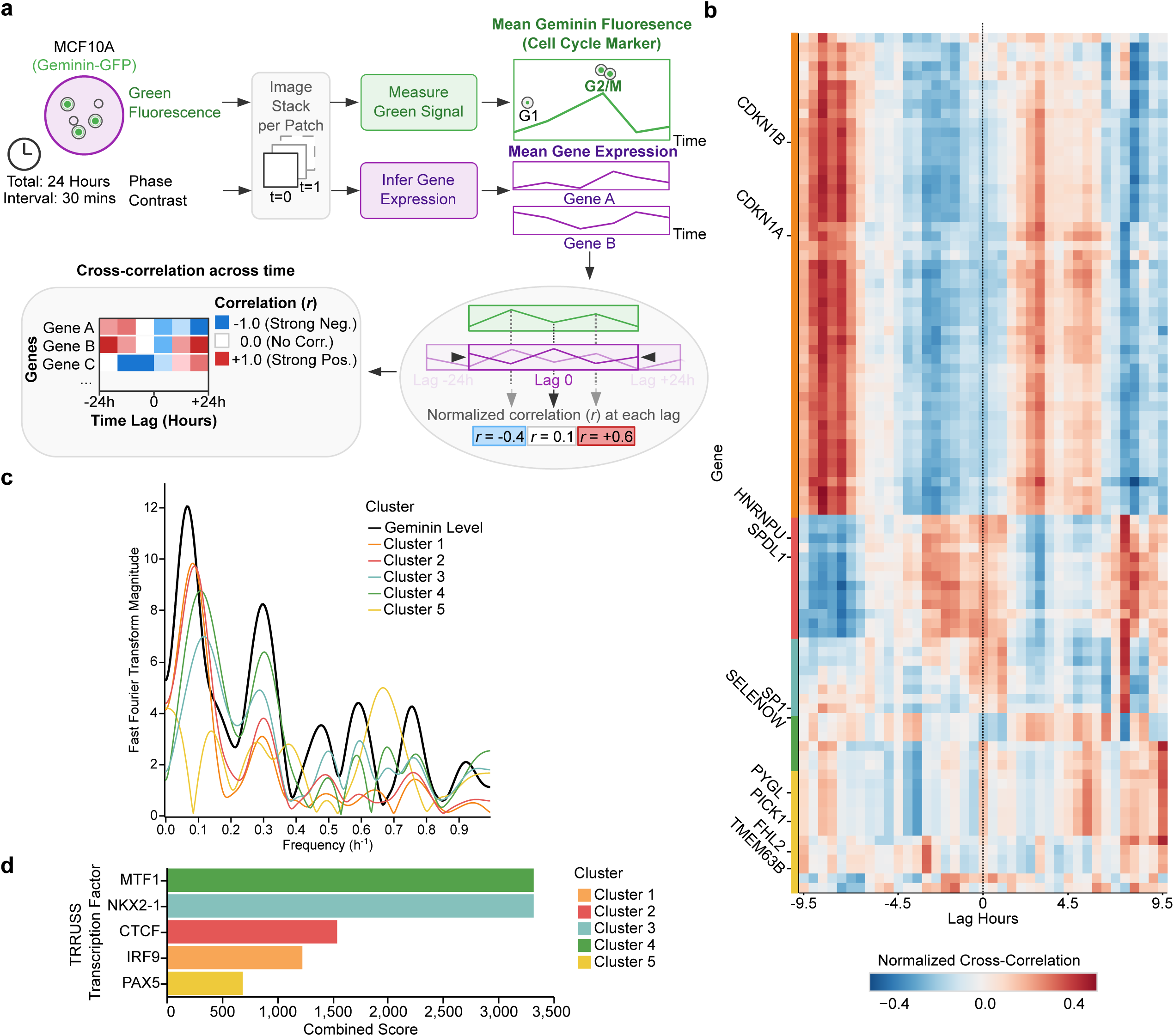
PENNE accurately tracks the temporal dynamics of gene-expression during mitosis. **a**, Workflow of the mitosis tracking experiment. We tracked the temporal dynamics of geminin along with PENNE-predicted gene-expression profiles across 24 hours with images taken every 30 minutes. We compared the cross-correlation between the predicted gene-expression profiles and the actual geminin levels across different time lags. **b**, Heatmap showing the normalized cross-correlation between the PENNE-predicted gene-expression levels and geminin fluorescence across different time lags for each of the high-confidence genes. We observed a periodic pattern of correlation for many genes, as well as five clusters of genes with similar temporal correlation patterns (left colourbar: cluster). The time lag of 0 (dashed line) is where the predicted gene-expression levels are directly compared to the actual geminin levels without any time shift. **c**, Fast Fourier transform analysis of the cross-correlation to identify the dominant frequencies, represented by the peaks, of the temporal correlation. We observed a dominant frequency of ≈0.05 cycles/h for both geminin and the correlation of each cluster, which corresponds to a period of ≈20 h, consistent with the expected cell-cycle duration for MCF10A cells. **d**, Enrichment analysis of the using the TRRUST database^104^ to identify the top-most enriched transcription factors within each cluster. Growth-related transcription factors such as CTCF and MTF1 are enriched in cluster 1, which has a positive correlation with geminin at lag 0, whereas cell-cycle-checkpoint-related transcription factors including IRF9 are enriched in cluster 2, which has a negative correlation with geminin at lag 0 but a positive correlation at a later time lag.

Having established that PENNE accurately captures canonical cell-cycle dynamics (clusters 1-3), we investigated clusters 4 and 5 to identify potentially uncharacterized temporal periodicities. We found that cluster 4 is highly enriched in MTF1, a transcription factor known to be involved in cell cycle regulation by sensing and responding to cellular stress.^74^ For example, MTF1 regulates the expression of *SELENOW*, a gene in the cluster 4, to mitigate oxidative stress during the G1 phase of the cell cycle in breast cancer cells.^75^ This function is consistent with the negative correlation of cluster 4 with geminin at lag 0, where geminin levels are low and cells are in G1 phase. However, we found that the cross-correlation fast Fourier transform cluster shows a periodicity of ≈10 h and 3 h, which is much shorter than the expected cell-cycle length. This finding suggests that stress-sensing genes in cluster 4 may be activated more frequently than the overall cell-cycle progression, potentially in response to the transient mechanical and metabolic stress of mitosis.^76^ Furthermore, there are also genes in cluster 4 whose expression is positively correlated with geminin levels at lag 0, suggesting that some stress response programs mediated by MTF1 may also be activated during G2/M phase, perhaps to mitigate the stress of actively dividing cells. By capturing this dynamic without external perturbation, our model generates a novel hypothesis for a plausible link between the biophysical and metabolic stress of mitosis and the activation of a stress response program mediated by MTF1.

Building on the observation of the transient physical changes during mitosis, further investigation of cluster 5 revealed a distinct temporal periodicity that highlights PENNE’s unique sensitivity to biophysical and metabolic shifts. Deeper inspection of the constituent genes revealed a highly coordinated network driving mitotic remodelling. As PENNE’s predictive architecture is grounded in phase-contrast imaging and therefore fundamentally captures physical density and gross morphological changes, our model identified a temporal gene cluster heavily enriched for mechanosensitive channels (*TMEM63B*),^77^ focal adhesion scaffolding (*FHL2*),^78^ membrane curvature regulators (*PICK1*),^79^ and rapid glycogen mobilization (*PYGL*)^80^ that peaks in synchrony with mitotic morphological changes to reconstruct the temporal dynamics of the cellular cytoskeleton and metabolome. Rather than capturing a strictly biochemical cell-cycle regulator that should have a similar periodicity to geminin, we hypothesize that cluster 5 represents the coupled responses of the cellular mechanosensitive and metabolic machinery to the transient stress of mitosis. This hypothesis is particularly interesting as there may be a role for mechanosensitive channel genes like *TMEM63B* in mitosis. Therefore, this finding opens up a new avenue for future research to investigate the role of this transient mitotic stress and biophysical response in regulating cell cycle progression. Overall, this application of PENNE highlights the model’s accuracy in inferring temporal cell dynamics and in enabling the discovery of novel gene-expression dynamics and hypothesis generation through continuous tracking of gene expression from live-cell PCM images.

## Discussion

While tracking temporal changes in gene expression is critical for understanding the dynamics of cellular states and behaviours, this objective is difficult to measure due to the destructive nature of sequencing. As such, current methods often rely on comparing different cells across independent samples captured at different time points, which at best introduces confounding factors and batch effects and at worst comparing different cells altogether. In this work, we developed PENNE, a high-resolution, transcriptome-wide method for inferring gene-expression profiles directly from live-cell images by leveraging the molecular knowledge learned from spatial transcriptomic H&E images through adversarial domain adaptation. We demonstrated that PENNE accurately infers cell-type-specific gene expression markers from PCM images, tracks the temporal dynamics of cellular confluency and radiation response, identifies a set of high-confidence genes whose expression can be reliably inferred from PCM images, and track cell-cycle progression to identify novel cell-cycle-related gene-expression dynamics. These findings suggest that PENNE enables new workflows orthogonal to traditional transcriptional methodologies that compare different cells across snapshots, and may spark new longitudinal approaches to discover novel therapeutic strategies and biological mechanisms.

Our findings suggest that the morphological features in images from non-stained modalities, such as PCM, contain rich information about the underlying gene-expression profiles of cells. As a result, future work can leverage similar architectures to create models that can infer gene-expression profiles from other non-stained and popular modalities such as bright-field,^81^ differential interference contrast,^82^ and Fourier ptychographic microscopies^83^ as those modalities may provide additional information of the cell morphology to improve the accuracy of inference. In addition, PENNE identified a set of genes with distinct temporal dynamics during cell-cycle progression. Future work can investigate the underlying mechanisms of these genes and identify both the role of MTF1 in mediating the transient stress response during mitosis and the role of mechanosensitive channels in mitosis. Understanding these mechanisms may provide new insights into the regulation of cell-cycle progression and unlock novel therapeutic strategies and targets for diseases, such as cancer, that involve dysregulation of the cell cycle.

Despite promising results and the potential novel workflow enabled by PENNE, there are still limitations that can be addressed in future work. While we observed successful one-shot transference to new human cell types and datasets not seen during training with PENNE, we used a training dataset that has a limited number of human cell types and morphological features. Future work can focus on training PENNE on a larger and more diverse dataset of both spatial transcriptomic H&E and PCM images of both human and non-human cell types to further improve the model generalizability and robustness. Additionally, while PENNE infers gene-expression profiles from live-cell PCM images, the resolution is still at patch level, which may not capture the full heterogeneity of gene expression at the single-cell level. Future work can further improve PENNE to infer gene-expression profiles at the single-cell level from PCM images by training a model using single-cell live-cell imaging data with matched single-cell RNA sequencing data or using a higher resolution modality such as Visium HD^84^ for training. This higher resolution of inferred gene expression at the single-cell level could provide more granular insights into the cellular heterogeneity and interaction of live-cell cultures. Finally, while we identified a set of high-confidence genes, future work could investigate the underlying mechanisms of how these genes are related to the morphological features in PCM images and how they contribute to the regulation of cellular states and behaviours. Overall, PENNE represents a significant advancement in the field of live-cell imaging by connecting the morphological features in PCM images to gene-expression profiles, and opens up new avenues for monitoring the temporal dynamics of cellular states and behaviours in a non-destructive and high-throughput manner.

## Methods

### PENNE model architecture overview

PENNE contains two major components: a pre-trained component for feature extraction and a trainable component for domain adaptation and gene-expression inference (Figure 1b). During training, the frozen pre-trained component extracts features from both spatial transcriptomic H&E-image patches and PCM-image patches, where we pass PCM patches through SPAGHETTI,^21^ a style transfer method that assists H&E-image-based models to generalize on PCM images by converting PCM images to H&E-like images. Through the SPAGHETTI framework, we pass H&E-like images to Phikon-v2,^16^ a foundation vision transformer based feature extractor intended for H&E images, which extracts features from both H&E patches and H&E-like PCM patches. We then passes the output features from Phikon-v2, which are vectors of length 1024 for both PCM and H&E, denoted as *f_p_* and *f_h_* respectively, to the trainable component of PENNE.

The trainable component of PENNE consists of five major modules, which are the Translator Module, the Domain Classifier Module, the Domain Discriminator Module, the Cell-Type Classifier Module, and the Gene-Expression Predictor Module. During training, the trainable component of PENNE learns the mapping from image features to gene-expression profiles from spatial transcriptomic H&E patches as well as the domain adaptation from PCM image features to H&E image features so that the model may apply the mapping to PCM image features despite the lack of ground-truth gene-expression data for PCM images. During inference, PENNE only uses the Translator Module and the Gene Expression Predictor Module to infer gene expression profiles from PCM image patches.

#### Translator Module

The Translator Module (*TM*) is a series of fully-connected layers that translates the output features from Phikon-v2 and separates the domain-specific information and the domain-invariant information into two orthogonal feature vectors. The output of this module is two vectors of lengths 64 and 960, representing the domain-invariant biological (*f_pb_* for PCM and *f_hb_* for H&E) and domain-specific features (*f_pd_* for PCM and *f_hd_* for H&E), respectively. To ensure that the domain-invariant features and the domain-specific features are independent, we apply an Independent Loss (*L_Independent_*) based on a combination of the Hilbert-Schmidt Independence Criterion (HSIC)^85^ and the cosine similarity between the domain-invariant and domain-specific features. The HSIC measures the statistical dependence between two sets of variables, where a value of zero indicates independence. The HSIC between two random variables *X* and *Y* is defined as

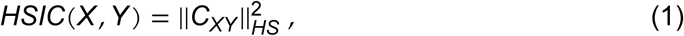

where *C_XY_* is the cross-covariance operator between the reproducing kernel Hilbert spaces (RKHS) of *X* and *Y*, and || ⋅ ||*_HS_* denotes the Hilbert-Schmidt norm. Here, we estimate the HSIC between the domain-invariant and domain-specific features for both PCM and H&E using a radial basis function (RBF) kernel^86^ for each batch of features during training:

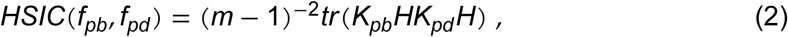

where *m* is the batch size, *K_pb_* and *K_pd_* are the kernel matrices defined as 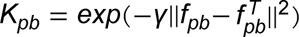 and 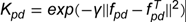 respectively, where *γ* is the kernel bandwidth parameter where we set to the median of the pairwise distances between the features in the batch. Furthermore, *H* is the centering matrix defined as 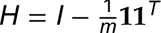 with identity matrix *I* and a vector of ones, **1**. Similarly, we compute *HSIC*(*f_hb_*, *f_hd_*) for all H&E features.

The cosine similarity loss component encourages orthogonality between the domain-invariant and domain-specific features, further promoting their independence. We first compute the orthogonality matrix by taking the dot product of the domain-invariant features with the transpose of the domain-specific features for both PCM and H&E features. The cosine similarity loss is then the squared L2 norm of the orthogonality matrix for both PCM and H&E features. Then, the loss for PCM features is

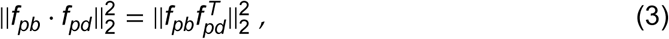

while the loss for H&E features is

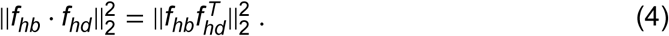

Finally, we define the overall Independent Loss as

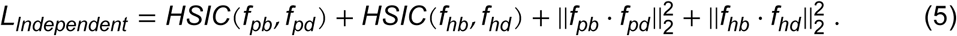

#### Domain Classifier Module

We pass the domain-specific features through the Domain Classifier Module (*DC*), which is a series of fully-connected layers that classifies whether the input features are from PCM or H&E images. This module ensures that the domain-specific features contain sufficient information to distinguish between the two domains and encourages the Translator Module to effectively separate domain-specific information from domain-invariant biological information. We train the Domain Classifier Module using a standard binary cross-entropy loss defined as:

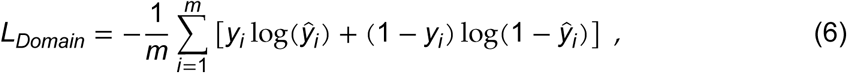

where *m* is the batch size, *y_i_* is the true label (1 for PCM, 0 for H&E) and *y^î^* is the predicted probability for the *i*-th sample in the batch.

#### Domain Discriminator Module

We pass the domain-invariant biological features through the Domain Discriminator Module (*DD*), which is another series of fully connected layers that aim to classify whether the input features are from PCM or H&E images and uses a standard binary cross-entropy loss, *L_Alignment_*, similar to Equation (6). However, unlike the Domain Classifier Module, we train the Domain Discriminator Module adversarially using a Gradient Reversal Layer (GRL) inspired by the Domain-Adversarial Neural Network (DANN) framework.^87^ The GRL inverts the gradients during backpropgation, encouraging the Translator Module to produce truly domain-invariant features that are indistinguishable between PCM and H&E domains. Therefore, we maximize *L_Alignment_* in order to better align the biological embedding between *f_pb_* and *f_hb_*.

In addition, to further ensure that the domain-invariant biological features of *f_pb_* and *f_hb_* are aligned in the feature space, we employ a Correlation Alignment loss (*L_Correlation_*) that minimizes the difference in the second-order statistics (covariances) between the two feature distributions.^88^ We define the Correlation Alignment loss as:

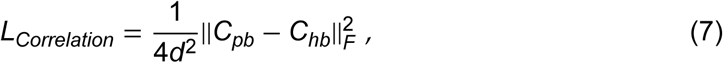

where *C_pb_* and *C_hb_* are the covariance matrices of the domain-invariant biological features from PCM and H&E, respectively, *d* is the dimensionality of the features (64 by default), and || ⋅ ||*_F_* denotes the Frobenius norm.

#### Cell-Type Classifier Module

Due to the lack of ground truth gene expressions for PCM images, we leverage cell-type annotations available for PCM images to further regularize the domain-invariant biological features and avoid overfitting to the spatial transcriptomic H&E data. We pass the domain-invariant biological features from PCM through the Cell-Type Classifier Module (*CC*), which is a series of fully-connected layers that classifies the cell type from input features. We train the Cell-Type Classifier Module using a standard multi-class cross-entropy loss, *L_Feature_*, to ensure that the domain-invariant biological features contain sufficient information to distinguish between different cell types in PCM as an additional biological constraint.

#### Gene-Expression Predictor Module

Finally, we pass the domain-invariant biological features from both PCM and H&E through the Gene-Expression Predictor Module (*GE*) to generate the predicted gene-expression profiles. The Gene-Expression Predictor Module uses a gated multi-layer perceptron architecture,^89^ which is effective in modelling long-range dependencies in high-dimensional data due to its use of spatial gating units as a self-attention mechanism. In PENNE, the added spatial gating units allow features from different dimensions to interact with each other, enabling the model to capture complex relationships between different genes.

To compute the gene-expression loss *L_Expression_*, due to the high dimensionality and sparsity of gene expression data, we use a combination of Huber loss^90^ and cosine loss from the predicted and ground-truth gene-expression profiles in H&E images to make the model more robust to outliers and noise in the data. The Huber loss is a combination of mean squared error (L2 loss) and mean absolute error (L1 loss), which is less sensitive to outliers than L2 loss alone:

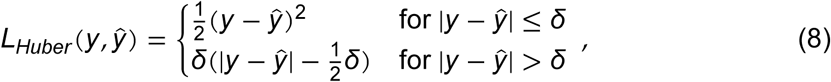

where *y* is the ground truth gene expression, *y*^ is the predicted gene expression, and *δ* is a hyperparameter that determines the threshold between L1 and L2 loss. Furthermore, the cosine loss measures the cosine similarity between the predicted and ground-truth gene-expression profiles, which helps to capture the overall direction of the gene-expression vectors rather than just their magnitude. Therefore, we define the overall expression loss as:

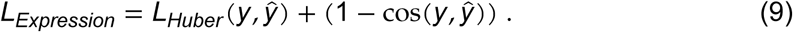

To enforce better generalization to PCM images, we obtained a list of marker genes for each cell type from the GDSC cell line dataset^91^ and computed an additional cross cell-type marker gene loss (*L_Marker_*) based on the margin ranking loss between the predicted expression levels of marker genes and non-marker genes for each cell type in the PCM images. We defined the margin ranking loss as:

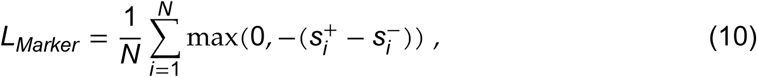

where *N* is the number of cell types, *s*^+^ is the predicted expression level of the marker gene for a cell type *i*, and *s*^−^ is the predicted expression level of the marker gene in other cell types.

Overall, to train the model, we used a combination of the above loss functions with various weights to balance the contributions of each component. The loss weights are tunable hyperparameters based on the specific dataset and task. We solve a minimax optimization problem to maximize *L_Alignment_* from *DD* while minimizing all other loss components to obtain the optimal parameters for PENNE:

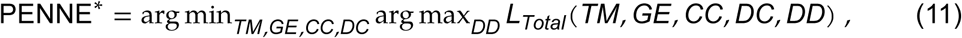

where *L_Total_* is the total loss function. *L_Total_* combines all the individual loss components with their respective weights:

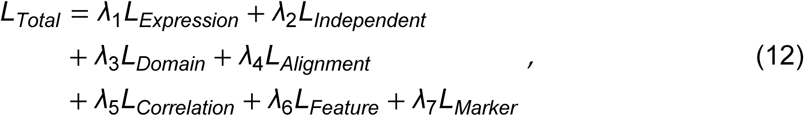

where *λ*_1_ to *λ*_7_ are loss-weight hyperparameters that control the relative importance of each loss component during training.

### Training details

We trained PENNE in three stages with a total of 46,161 paired H&E patches (size 224 px×224 px) and spatial transcriptomics data of 18,085 genes across all nine publicly-available human 10x Genomics Visium samples^27^ with available H&E images. For each Visium high-resolution image, we randomly cropped one of the 2,743 PCM images (original size 1,408 px×1,040 px) across six different cell types and confluencies from the training portion of LIVECell^28^ to be 224 px×224 px for domain adaptation.

During the first stage of training, we trained with a batch size of 16 for 20 epochs with the main objective to learn the mapping from H&E-image features to gene-expression profiles. As such, we set the loss weights to *λ*_1_ = 5.0 for *L_Expression_*, *λ*_2_ = 0.1 for *L_Independent_*, *λ*_3_ = 0.1 for *L_Domain_*, *λ*_4_ = 0.1 for *L_Alignment_*, *λ*_5_ = 0.1 for *L_Correlation_*, and all other loss weights to zero to encourage the model to learn the mapping from H&E features to gene-expression profiles while also regularizing the domain-invariant biological features with cell type information.

During the second stage, we trained with a batch size of 8 for 30 epochs with the main objective to learning the domain adaptation from PCM-image features to H&E-image features. Therefore, we increased the loss weights to *λ*_2_ = 3.0 for *L_Independent_*, *λ*_3_ = 10.0 for *L_Domain_*, *λ*_4_ = 3.0 for *L_Alignment_*, *λ*_5_ = 3.0 for *L_Correlation_*, *λ*_6_ = 8.0 for *L_Feature_*, and *λ*_7_ = 3.0 for *L_Marker_* to encourage the model to learn the domain adaptation while still maintaining some focus on learning the mapping from H&E features to gene-expression profiles.

During the third and final stage, we trained with a batch size of 8 for 30 epochs with the main objective to fine-tune the model for better gene expression inference on PCM images. Therefore, we increased the loss weight for *L_Marker_* to *λ*_7_ = 10.0 while keeping the other loss weights the same as the second stage to encourage the model to generate more accurate gene-expression profiles for PCM images by leveraging the cell-type-specific marker-gene information as a strong biological constraint.

We trained the model with an 80% of the data and tested with the remaining 20%. We trained using an AdamW optimizer^92^ with a learning rate initially at 0.005 with a step decay Gamma of 0.1 every 10 epochs.

### Cell culture, imaging, and bulk RNA sequencing and preprocessing

We first established a validation dataset using MCF10A and HCT116 cell lines. We cultured H2B-GFP-expressing MCF10A cells (ATCC, CRL-10317) in 1:1 mixture of F12:DMEM media supplemented with 5% horse serum (Wisent Bioproducts, 098150), 20 ng/mL human EGF (Cedarlane Labs, AF-100-15), 0.5 mg/mL hydrocortisone (Sigma-Aldrich, H0888), 100 ng/mL cholera toxin (Sigma-Aldrich, C8052), and 10 mg/mL recombinant human insulin (SAFC, 91077C). We also cultured unlabeled HCT116 p53-null cells in McCoy’s 5A medium, and 10% fetal bovine serum. All media were supplemented with 1% penicillin-streptomycin. We then analyzed H2B-GFP-expressing MCF10A cells and unlabeled HCT116 p53-null cells both as monocultures and as a 50-50 mixed co-culture under non-irradiated (NIR) and 10 Gy irradiated (IR) conditions, with three biological replicates per group. We monitored the cells by live-cell imaging on a Sartorius Incucyte SX5 Live Cell Analysis System, taking both phase-contrast and green fluorescence images every 24 hours to acquire images over a 72-hour period.

We harvested the total RNA at 72 hours post-irradiation using TriZol reagent (Life Technologies, 15596018) for poly(A)-selected mRNA-seq using NovaSeq X Plus paired-end 150-bp sequencing (Novogene). We obtained an average of 20 million reads per sample across 18 samples (3 replicates for each of the 6 conditions: MCF10A NIR, MCF10A IR, HCT116 NIR, HCT116 IR, 1:1 MCF10A:HCT116 NIR, and 1:1 MCF10A:HCT116 IR). To preprocess the bulk RNA sequencing data and align with genomic data, we first performed quality control and trimmed the adapters from the raw RNA sequencing reads using fastp.^93^ We then generated a genome index using the human reference genome (GRCh38) and gene annotation files from GENCODE^94^ and aligned the processed reads using the STAR aligner.^95^ Finally, we quantified the gene expression levels using featureCounts^96^ to obtain the raw gene counts for each sample, which were then log-counts-per-million normalized to be used for downstream analysis and comparison with the predicted gene expression profiles from PENNE.

To establish the cell-culture images for mitosis tracking, we built a live-cell imaging assay using the untreated and 5 Gy irradiated MCF10A human breast epithelial cells (CRL-10317, ATCC). We tagged a green fluorescent protein, Clover, to a partial gene sequence of geminin (amino acids 1-110), namely the localization sequence, through lentiviral transduction (Addgene, plasmid no. 83915). We grew cells in 1:1 mixture of F12:DMEM media supplemented with 5% horse serum (Wisent Bioproducts, 098150), 20 ng/mL human EGF (Cedarlane Labs, AF-100-15), 0.5 mg/mL hydrocortisone (Sigma-Aldrich, H0888), 100 ng/mL cholera toxin (Sigma-Aldrich, C8052), 10 mg/mL recombinant human insulin (Sigma-Aldrich, 91077C), and 1% penicillin-streptomycin (Fisher Scientific, 15140122). We used Sartorius Incucyte S3 Live Cell Analysis System to acquire images for longitudinally tracking single cells and took both phase-contrast and green fluorescence images every 30 minutes for 7 days. We only analyzed the images for the first 24 hours to avoid over confluency.

### Cell-type marker gene-expression analysis

We obtained a list of marker genes for each cell type from the GDSC cell-line dataset^91^ and GeneRIF,^97^ a database that provides gene-function annotations based on literature mining. We obtained the mean log-counts-per-million normalized expression level of the cell-type marker genes from the predicted gene-expression profiles across all testing 7,104 LIVECell PCM image patches and 540 U373 cell-line images.^32^ We also performed random permutation of the input images as a negative control before passing the images into PENNE for inferences. We compared the statistical significance of the mean expression levels of these marker genes across the predicted profiles using Mann-Whitney *U* tests. For each cell type, we also performed differentially expressed gene (DEG) analysis and gene set enrichment analysis (GSEA)^35^ using the PreRank method from GSEApy^98^ to test whether the marker genes for that cell type are significantly enriched for that cell type.

Furthermore, we generated gene expression inferences from PCM images for a 1:1 mixture of MCF10A and HCT116 cells. We also measured the greenness intensity, a measure for the amount of MCF10A cells in the image, for each image patch by taking the percentage of green pixels above a dynamic background threshold over the total cell area. We then built a ridge regression model to predict the greenness intensity from the predicted greenness intensity from the predicted gene expression. We also compute the Spearman’s correlation between the 10% most positively-correlated genes and most negatively-correlated genes and performed EnrichR^99^ on the MSigDB Hallmark dataset^100^ using GSEApy.

### Confluency and psuedo-time analysis

We generated the gene expression prediction with PENNE for 2,592 PCM images of pure MCF10A cells, pure HCT116 cells, and a 1:1 mixture of MCF10A and HCT116 cells at three different confluencies (imaged at day 0, day 1, and day 2 after seeding). We then computed the pseudo-time ordering of these images based on the predicted gene-expression profiles using Monocle3^101^ by setting a random image at day 0 as the root. To obtain a list of programs that are enriched in day 2 compared to day 0, we performed DEG analysis followed by PreRank GSEA using the KEGG database.^102^ We also computed the mean expression levels of all genes in the MSigDB Hallmark Apical Junction gene set using inferences from both original and permuted PCM images as input to PENNE. We compared the statistical significances of the mean expression levels using Mann-Whitney *U* tests.

### Radiation response analysis

We inferred the gene expressions for pure MCF10A cells, pure HCT116 cells, and a 1:1 mixture of MCF10A and HCT116 cells at different confluencies (day 0, day 1, and day 2 after seeding) and different treatments (with either 0 Gy or 10 Gy radiation). We then clustered and visualized gene-expression profiles of images using TooManyCells^55^ and TooManyCellsInteractive,^56^ respectively. We then performed DEG analysis followed by EnrichR using the most differentially expressed genes between day 2 and day 0 for both pure MCF10A and pure HCT116 cells to identify the top enriched radiation damage-related terms in the MSigDB Hallmark database.

### High-confidence gene identification and model ablation study

We computed the mean Spearman’s correlation and explained variance between the pseudo-bulk predicted gene expression profiles from PCM images and the bulk RNA sequencing data for each gene across all samples. We then defined the high-confidence genes as the genes with a mean Spearman’s correlation greater than 0.3 and an explained variance greater than 0.5, where the explained variance between the predicted (*y*^) and ground truth gene expression (*y*) is defined as:

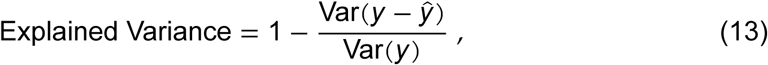

where 1 represents the maximum possible explained variance where the predicted gene expression perfectly captures the variance of the ground-truth gene expression. We then performed EnrichR on the Gene Ontology (GO) Biological Process database^103^ using the high-confidence genes to identify the top enriched biological processes that are captured by PENNE’s predictions. We also computed the sample correlation between the ground truth and predicted gene expression profiles using the high-confidence genes.

We performed an ablation study by removing different combination of the loss components during training and comparing the sample correlation between the ground-truth and predicted gene-expression profiles using all genes and the high-confidence genes to evaluate the importance of each loss component for the overall performance of PENNE.

### Gating and interpretability analysis

We extracted the gating values from the spatial gating units in the Gene Expression Predictor Module of PENNE for each high confidence predicted gene expression profile from PCM images. We then computed the mean gating values across all samples for each gene and plotted a Spearman’s correlation heatmap between the mean gating values and the predicted expression levels of each gene across all samples. We then performed hierarchical clustering to identify clusters and performed EnrichR on GO Cellular Component database using the genes in each cluster. We repeated this process by inputting randomly permuted PCM images.

### Longitudinal gene-expression inference across mitosis

We generated gene-expression predictions of all high confidence genes from PENNE for a time-lapse PCM imaging dataset of patches of MCF10A cells with a G2/M cell cycle marker (Geminin-GFP) across 24 hours with images taken every 30 minutes for 24 hours. We also quantify the level of geminin using an integrated intensity. We then computed an average of the gene expression levels and geminin levels for each time point across all patches. We conducted a cross-correlation analysis by shifting the predicted expression levels of the G2/M marker gene across different time lags and computing the normalized cross-correlation between the shifted predicted gene expression levels and the actual geminin levels, where −1 represents perfect negative correlation and 1 represents perfect positive correlation. We plotted a cross-correlation heatmap using time points up to 10 hours to avoid edge effects and performed hierarchical clustering to identify clusters of genes with similar temporal correlation with geminin. We performed a fast Fourier transform^63^ on the correlation to identify the dominant frequencies of the temporal correlation, as well as conducted EnrichR on the TRRUST database^104^ to identify the top enriched transcription factors for the genes in each cluster.

### Differential gene expression and enrichment analysis

We computed log fold changes using an ordinary least squares regression model and significance of the fold changes using a *t*-test. We reported the top enriched terms with an adjusted *p* < 0.250 after Benjamini-Hochberg correction for multiple hypothesis testing for both negative and positive enrichment. We reported normalized enrichment score (NES) for PreRank GSEA using all genes. We also reported the combined scores for enrichment from EnrichR using the most significantly differentially-expressed genes by choosing genes with a log fold change > 1 and adjusted *p* < 0.050.

## Data Availability

The spatial transcriptomics data used to train PENNE are publicly available from the 10x Genomics Visium Spatial Gene Expression Dataset.^27^ The live-cell PCM images used for domain adaptation and validation are publicly available from the LIVECell dataset.^28^ The U373 dataset used for validation of cell type marker can be obtained from the original publication.^32^

The bulk RNA sequencing data generated in this study have been deposited in the Gene Expression Omnibus (GEO) under accession number GSE338040. The imaging data generated in this study are available from Zenodo.^105^

## Code Availability

PENNE is open source and freely available at https://github.com/schwartzlab-methods/penne. The code for all analyses and figure generation is available at https://github.com/schwartzlab-methods/penne_paper_figures.

## Acknowledgments

We would like to thank the Vector Institute for Artificial Intelligence for providing the computing infrastructure and resources for this work.

## Authors Contributions

W. S. conceived the project. G. W. S. and C. M. supervised the project. S. M., M. M. T., and S. M. H. developed the dataset for validation. Z. F. D. developed the PENNE method, software, and benchmarks. Z. F. D. ran and analyzed benchmarks. Z. F. D. generated experimental results. Z. F. D. ran and analyzed data. Z. F. D., M. M. T., S. M., and G. W. S. wrote and edited the manuscript. All authors reviewed the manuscript.

