## Supplemental Information for "Longitudinal whole transcriptomic profiling of live cells through domain adaptation"

### Supplementary Information

#### Supplementary Notes

##### Supplementary Note S1: A complete list of all high-confidence genes

*ADGRG4*
*AIP*
*ARHGDIA*
*ASS1*
*ATG101*
*BIRC6*
*BRD8*
*CALM3*
*CAMK2N1*
*CAND1*
*CDKN1A*
*CDKN1B*
*CLASRP*
*CLK2*
*CNOT9*
*CSTF3*
*CUL3*
*DAZAP1*
*DDX23*
*DEK*
*DHX15*
*DUT*
*ENY2*
*FHL2*
*FKBP3*
*GMPR2*
*GMPS*
*HNRNPD*
*HNRNPH3*
*HNRNPL*
*HNRNPU*
*HNRNPUL1*
*ID1*
*ILF3*
*IMPA2*
*ITPK1*
*KIF22*
*KLHDC3*
*LAMP1*
*MBNL1*
*MCM7*
*MELK*
*NABP2*
*NDUFB6*
*NOP56*
*NSL1*
*NSUN2*
*NT5C*
*NUP107*
*PAGR1*
*PICK1*
*PIP4K2C*
*PLPP6*
*POLH*
*PREPL*
*PSAP*
*PSKH1*
*PTMS*
*PYGL*
*QDPR*
*RAD21*
*RAD23A*
*RIMS4*
*RSRC2*
*SART1*
*SELENOW*
*SLC27A4*
*SLC44A1*
*SMARCA4*
*SMARCB1*
*SMC1A*
*SNRPA*
*SP1*
*SPDL1*
*SRSF2*
*SRSF4*
*SRSF7*
*SSRP1*
*SUN2*
*TAP1*
*TMEM63B*
*TOP2B*
*TPR*
*TRAK1*
*TTYH1*
*U2SURP*
*UACA*
*UBALD2*
*URB1*
*VAT1L*
*WDR82*
*ZNF696*

#### Supplementary Figures

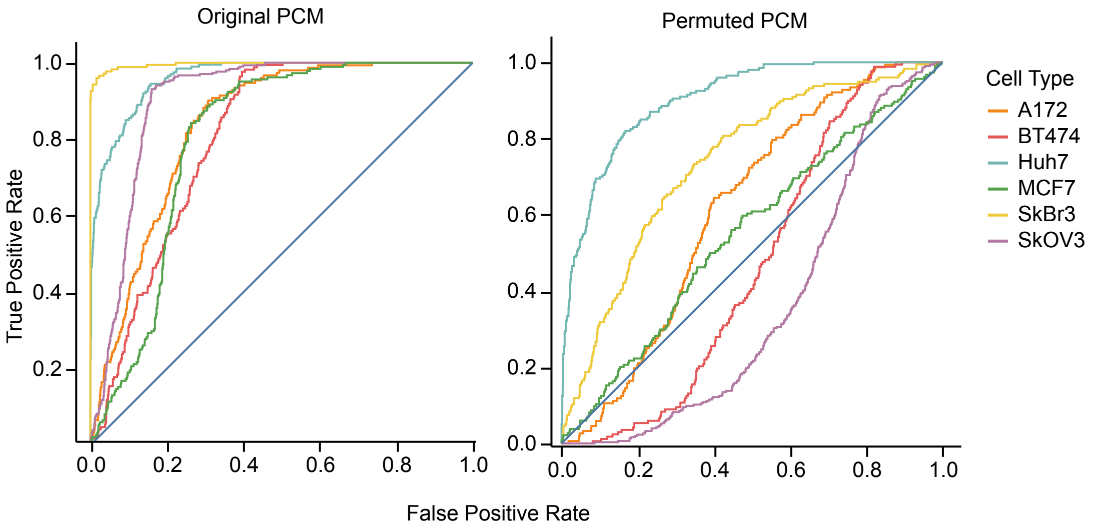

Original PCM
Permuted PCM
True Positive Rate
False Positive Rate
0.0
0.2
0.4
0.6
0.8
1.0
Cell Type
A172
BT474
Huh7
MCF7
SkBr3
SkOV3

Supplementary Figure S1: Area under the receiver operating characteristic (AUROC) curves for predicting the correct cell type using marker gene expression profiles for each cell type in the LIVECell dataset, using the original live-cell PCM images (left) and randomly permuted PCM images (right). The AUROC curves for the original PCM images are higher than those for the randomly permuted PCM images across all cell types, suggesting that PENNE can capture cell-type-specific gene-expression patterns from PCM images through morphological features. Blue line presents the random classifier baseline of an AUROC of 0.5.

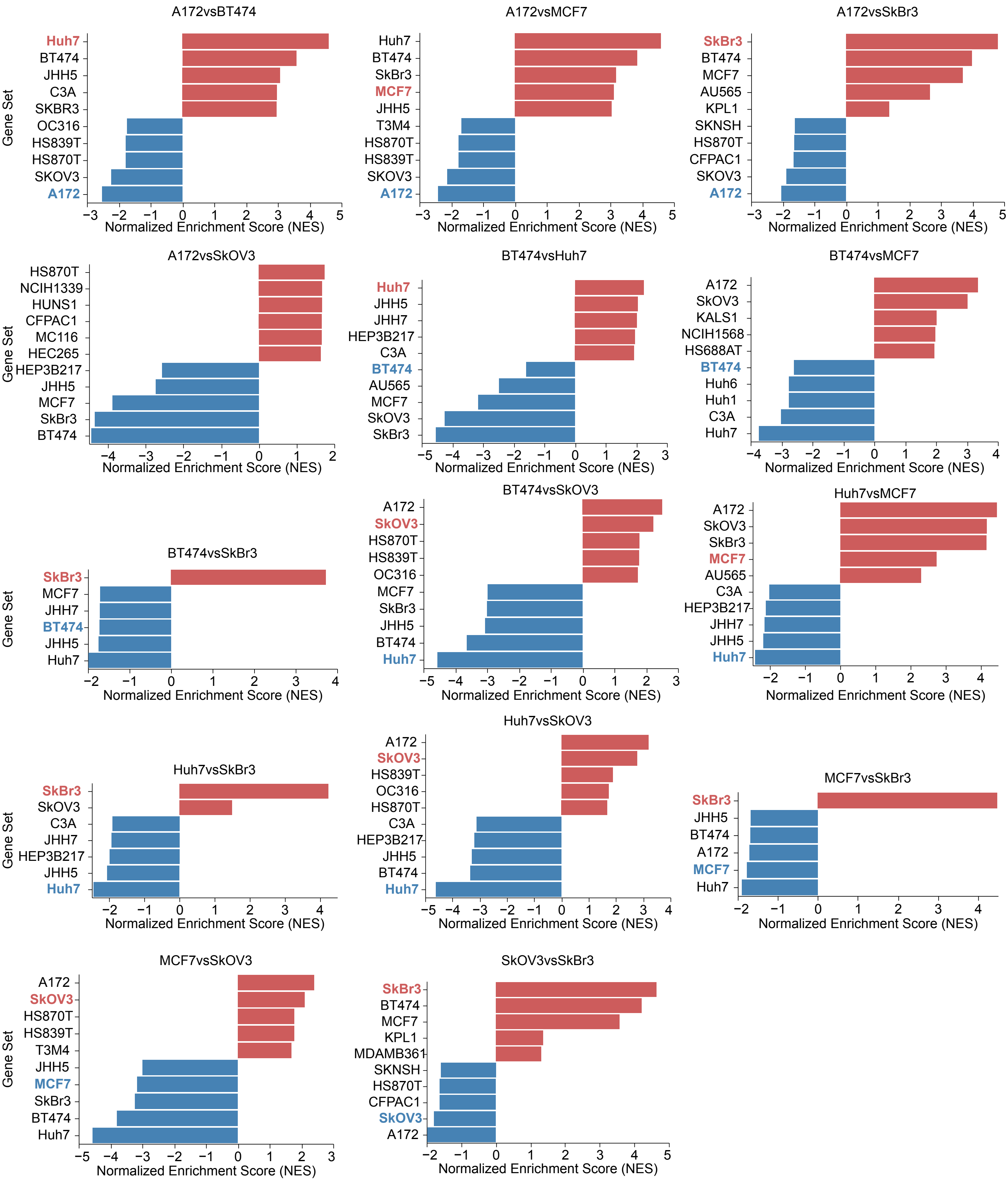
A172vsBT474
Huh7
BT474
JHH5
C3A
SKBR3
OC316
HS839T
HS870T
SKOV3
A172
A172vsMCF7
Huh7
BT474
SkBr3
MCF7
JHH5
T3M4
HS870T
HS839T
SKOV3
A172
A172vsSkBr3
SkBr3
BT474
MCF7
AU565
KPL1
SKNSH
HS870T
CFPAC1
SKOV3
A172
A172vsSkOV3
HS870T
NCIH1339
HUNS1
CFPAC1
MC116
HEC265
HEP3B217
JHH5
MCF7
SkBr3
BT474
BT474vsHuh7
Huh7
JHH5
JHH7
HEP3B217
C3A
BT474
AU565
MCF7
SkOV3
SkBr3
BT474vsMCF7
A172
SkOV3
KALS1
NCIH1568
HS688AT
BT474
Huh6
Huh1
C3A
Huh7
BT474vsSkBr3
SkBr3
MCF7
JHH7
BT474
JHH5
Huh7
BT474vsSkOV3
A172
SkOV3
HS870T
HS839T
OC316
MCF7
SkBr3
JHH5
BT474
Huh7
Huh7vsMCF7
A172
SkOV3
SkBr3
MCF7
AU565
C3A
HEP3B217
JHH7
JHH5
Huh7
Huh7vsSkBr3
SkBr3
SkOV3
C3A
JHH7
HEP3B217
JHH5
Huh7
Huh7vsSkOV3
A172
SkOV3
HS839T
OC316
HS870T
C3A
HEP3B217
JHH5
BT474
Huh7
MCF7vsSkBr3
SkBr3
JHH5
BT474
A172
MCF7
Huh7
MCF7vsSkOV3
A172
SkOV3
HS870T
HS839T
T3M4
JHH5
MCF7
SkBr3
BT474
Huh7
SkOV3vsSkBr3
SkBr3
BT474
MCF7
KPL1
MDAMB361
SKNSH
HS870T
CFPAC1
SkOV3
A172
Gene Set
Normalized Enrichment Score (NES)

Supplementary Figure S2: GSEA normalized enrichment scores showing the top 5 significant and most-enriched cell-type-related gene sets[91] from the differentially expressed genes between each pair of cell type in the LIVECell dataset. 12 out of 14 combinations show significant enrichment in the matched cell type, where the expected enrichment is highlighted for reference (blue) and compared (red) cell types.

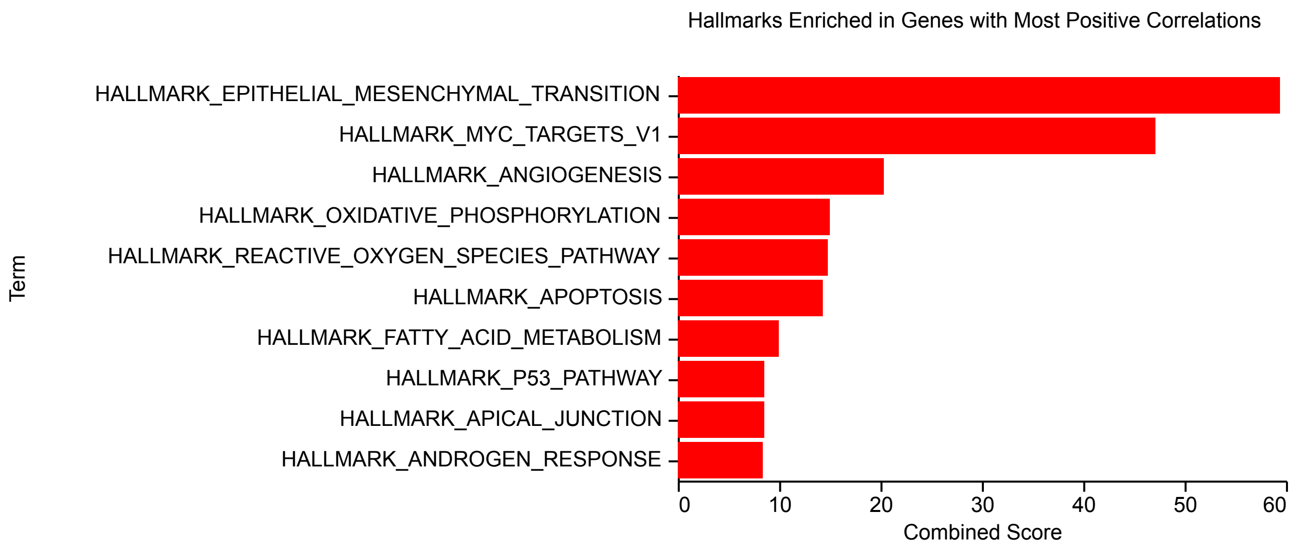

Hallmarks Enriched in Genes with Most Positive Correlations
HALLMARK_EPITHELIAL_MESENCHYMAL_TRANSITION
HALLMARK_MYC_TARGETS_V1
HALLMARK_ANGIOGENESIS
HALLMARK_OXIDATIVE_PHOSPHORYLATION
HALLMARK_REACTIVE_OXYGEN_SPECIES_PATHWAY
HALLMARK_APOPTOSIS
HALLMARK_FATTY_ACID_METABOLISM
HALLMARK_P53_PATHWAY
HALLMARK_APICAL_JUNCTION
HALLMARK_ANDROGEN_RESPONSE
Term
0
10
20
30
40
50
60
Combined Score


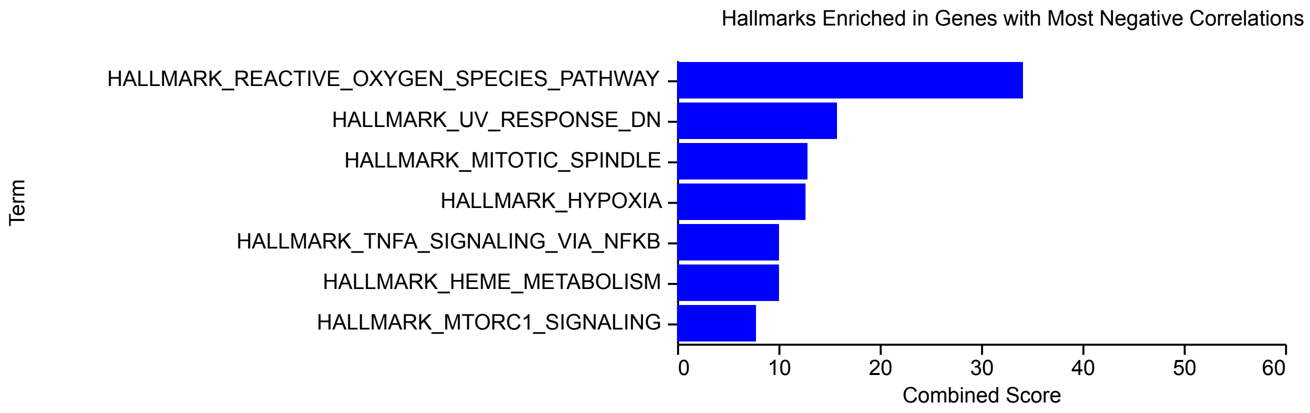

Hallmarks Enriched in Genes with Most Negative Correlations
HALLMARK_REACTIVE_OXYGEN_SPECIES_PATHWAY
HALLMARK_UV_RESPONSE_DN
HALLMARK_MITOTIC_SPINDLE
HALLMARK_HYPOXIA
HALLMARK_TNFA_SIGNALING_VIA_NFKB
HALLMARK_HEME_METABOLISM
HALLMARK_MTORC1_SIGNALING
Term
0
10
20
30
40
50
60
Combined Score

Supplementary Figure S3: Enrichment analysis of the most positively (top) and negatively (bottom) significantly-correlated genes from the MCF10A/HCT116 co-culture experiment using the MSigDB Hallmark dataset.[100] The most positively-correlated genes are significantly enriched in pathways related to cellular integrity while the most negatively-correlated genes are significantly-enriched for proliferation-related pathways.

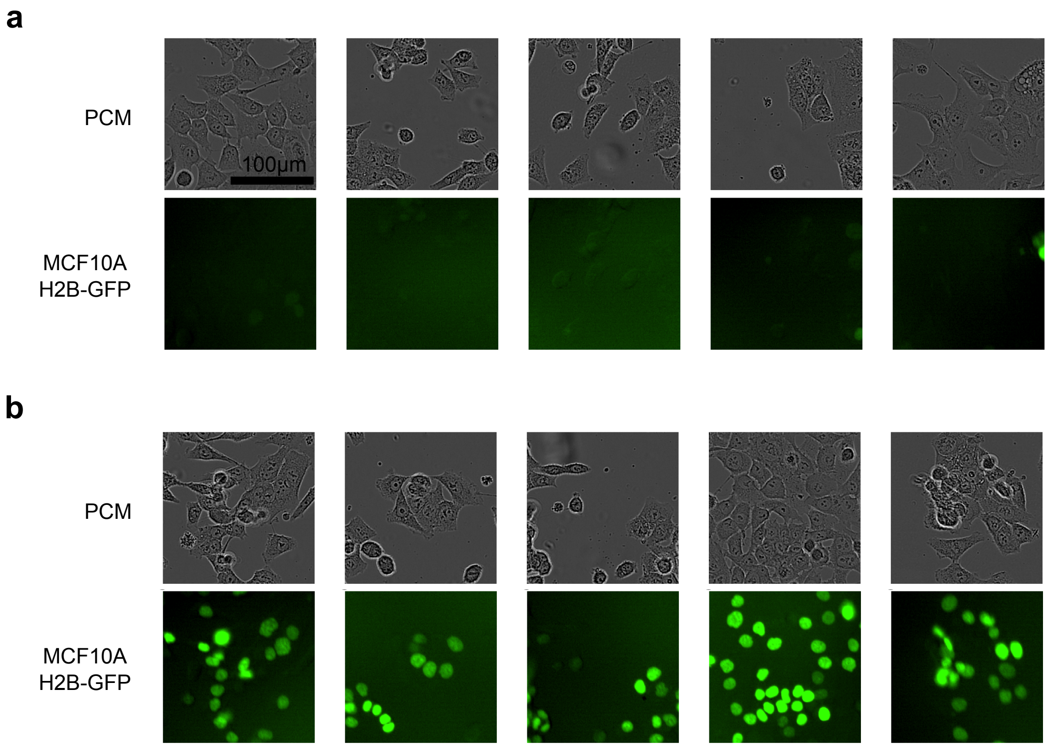
a
PCM
100µm
MCF10A
H2B-GFP
b
PCM
MCF10A
H2B-GFP

Supplementary Figure S4: Examples of PCM (top) and green fluorescence channel (bottom) GFP signal images in the MCF10A/HCT116 co-culture experiment. As MCF10A cells are tagged with H2B-GFP, the GFP signal serves as a measure of the fraction of MCF10A cells in each image. **a**, **b**, Examples of low GFP signal patches with a low fraction of MCF10A cells and a high fraction of HCT116 cells (**a**) or vice versa (**b**).

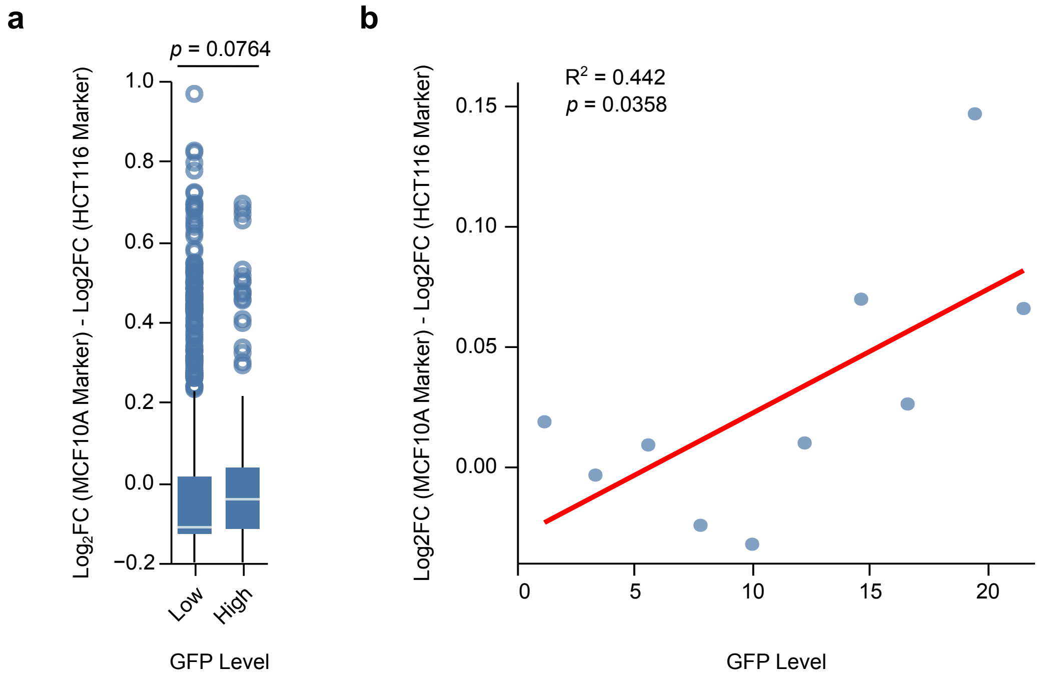
a
b
p = 0.0764
Log2FC (MCF10A Marker) - Log2FC (HCT116 Marker)
1.0
0.8
0.6
0.4
0.2
0.0
−0.2
Low
High
GFP Level
R2 = 0.442
p = 0.0358
0.15
0.10
0.05
0.00
0
5
10
15
20
Log2FC (MCF10A Marker) - Log2FC (HCT116 Marker)
GFP Level

Supplementary Figure S5: Difference in predicted expression of marker genes between MCF10A and HCT116 cells. **a**, Box-and-whisker plot showing the difference in predicted MCF10A marker gene expression and HCT116 marker gene expressions for high GFP signal patches (high MCF10A fraction) compared to low GFP signal patches (low MCF10A fraction). Statistical significance was determined using a two-tailed Wilcoxon rank-sum test ($p = 0.0764$). **b**, Scatter plot and regression line showing the difference in predicted expression of MCF10A and HCT116 marker genes compared to the GFP signal (MCF10A fraction) for each level of GFP. Statistical significance was determined using an $F$ test ($p = 0.0358$).

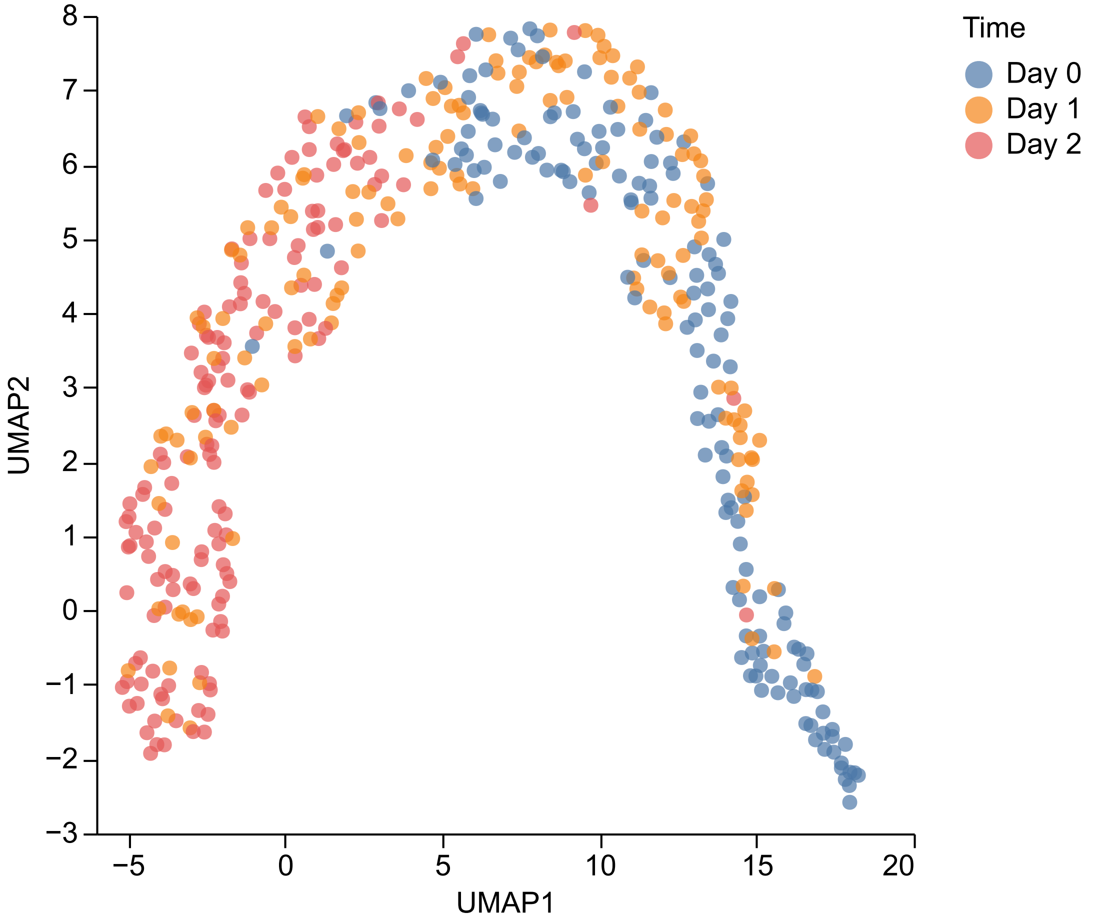

Time
Day 0
Day 1
Day 2
UMAP1
UMAP2
−5
0
5
10
15
20
−3
−2
−1
0
1
2
3
4
5
6
7
8

Supplementary Figure S6: UMAP projection of the PENNE-predicted gene-expression profiles for images at day 0 (least confluent), day 1 (moderately confluent), and day 2 (most confluent). The predicted gene-expression profiles show a clear separation between day 0 and day 2, with day 1 images showing an intermediate profile.

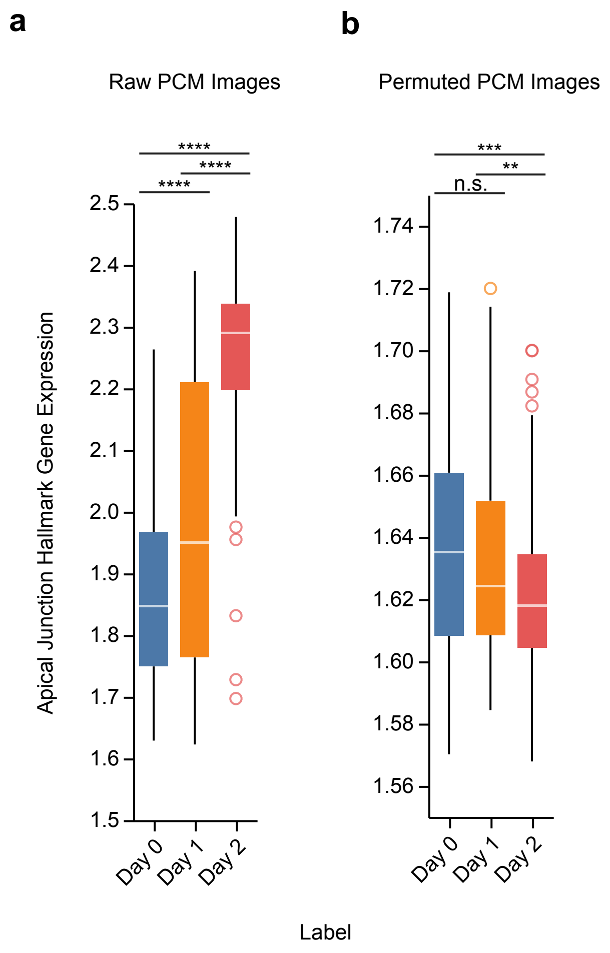
a
b
Raw PCM Images
Permuted PCM Images
****
****
****
***
**
n.s.
Apical Junction Hallmark Gene Expression
2.5
2.4
2.3
2.2
2.1
2.0
1.9
1.8
1.7
1.6
1.5
1.74
1.72
1.70
1.68
1.66
1.64
1.62
1.60
1.58
1.56
Day 0
Day 1
Day 2
Label

Supplementary Figure S7: **a**, **b**, Box-and-whisker plot showing the difference in predicted expression of the MSigDB Hallmark Apical Junction gene set for day 0, day 1, and day 2 images for both original (**a**) and permuted (**b**) PCM images. We observed a significant increasing trend as confluency increases as expected in the original PCM images but a decreasing trend in the permuted PCM images. Two-tailed Mann-Whitney *U* test, n.s.: not significant, ****: $p < 1.00 \times 10^{-4}$, ***: $p < 1.00 \times 10^{-3}$, **: $p < 0.0100$.

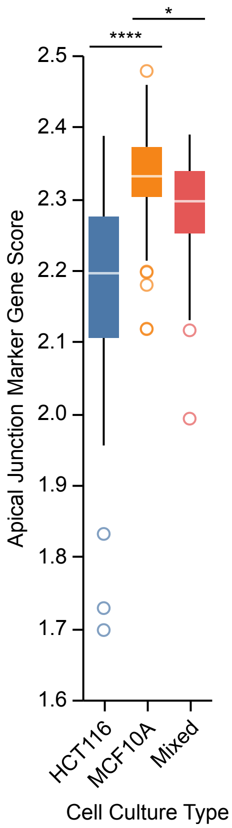

*
****
2.5
2.4
2.3
2.2
2.1
2.0
1.9
1.8
1.7
1.6
Apical Junction Marker Gene Score
HCT116
MCF10A
Mixed
Cell Culture Type

Supplementary Figure S8: Box-and-whisker plot showing the difference in PENNE-predicted expression of the MSigDB Hallmark Apical Junction gene set for day 2 images for pure MCF10A, pure HCT116, and mixed MCF10A/HCT116 cell-type live-cell images. We found a significantly higher expression of the Apical Junction gene set in pure MCF10A images compared to both pure HCT116 images and mixed MCF10A/HCT116 images, which is consistent with the expected higher expression of apical junction-related genes in epithelial cells like MCF10A compared to cancer cells such as HCT116. Two-tailed Wilcoxon rank-sum test, ****: $p < 1.00 \times 10^{-4}$, **: $p < 0.0500$.

**a**

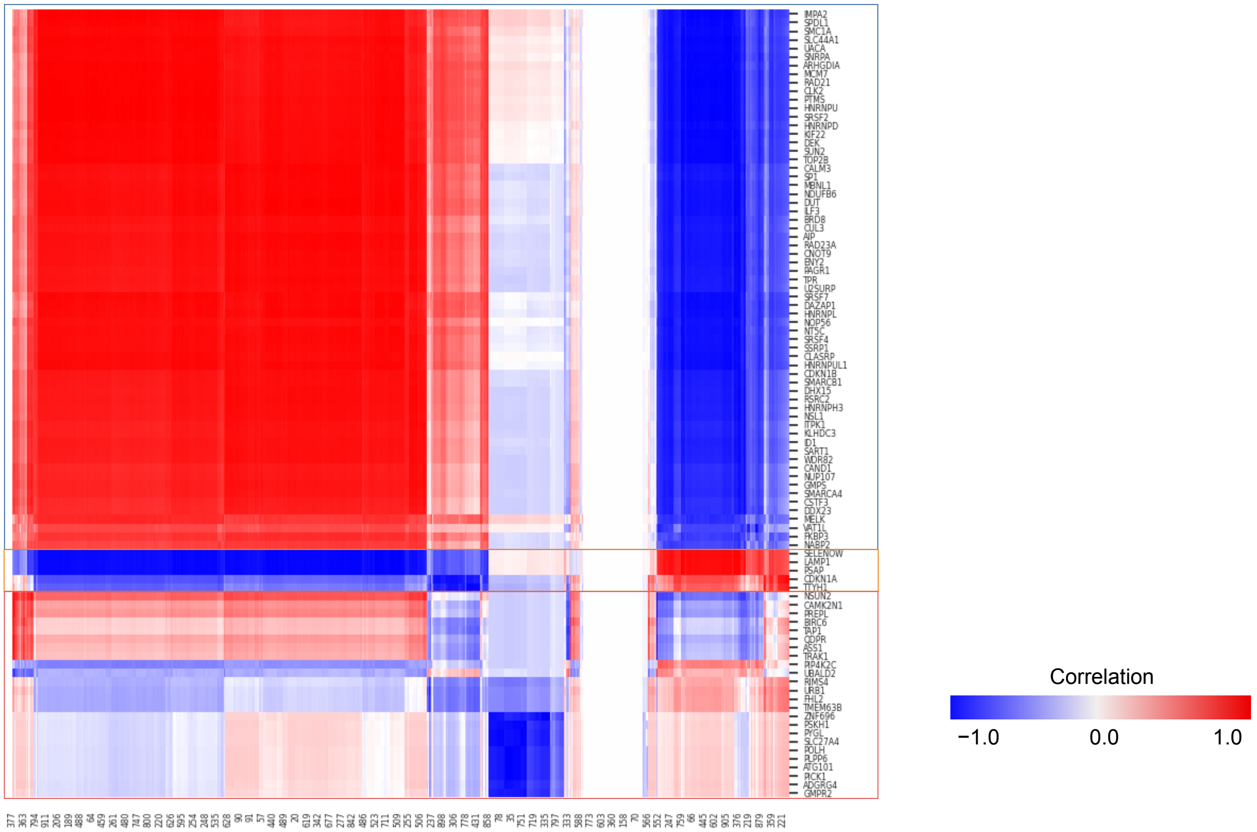
Correlation
−1.0
0.0
1.0

**b**

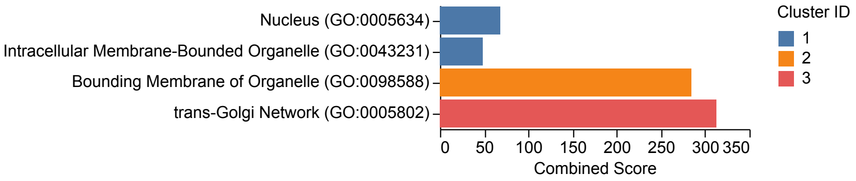
Nucleus (GO:0005634)
Intracellular Membrane-Bounded Organelle (GO:0043231)
Bounding Membrane of Organelle (GO:0098588)
trans-Golgi Network (GO:0005802)
0
50
100
150
200
250
300
350
Combined Score
Cluster ID
1
2
3

Supplementary Figure S9: PENNE's gating units capture organelle specific gene programs. **a**, Spearman's correlation heatmap between the mean gating values from the spatial gating units in the Gene Expression Predictor Module of PENNE and the predicted expression levels of each high confidence gene across all patches. **b**, Enrichment analysis of each cluster in **a** showing that each cluster is significantly enriched for genes related to specific organelles.

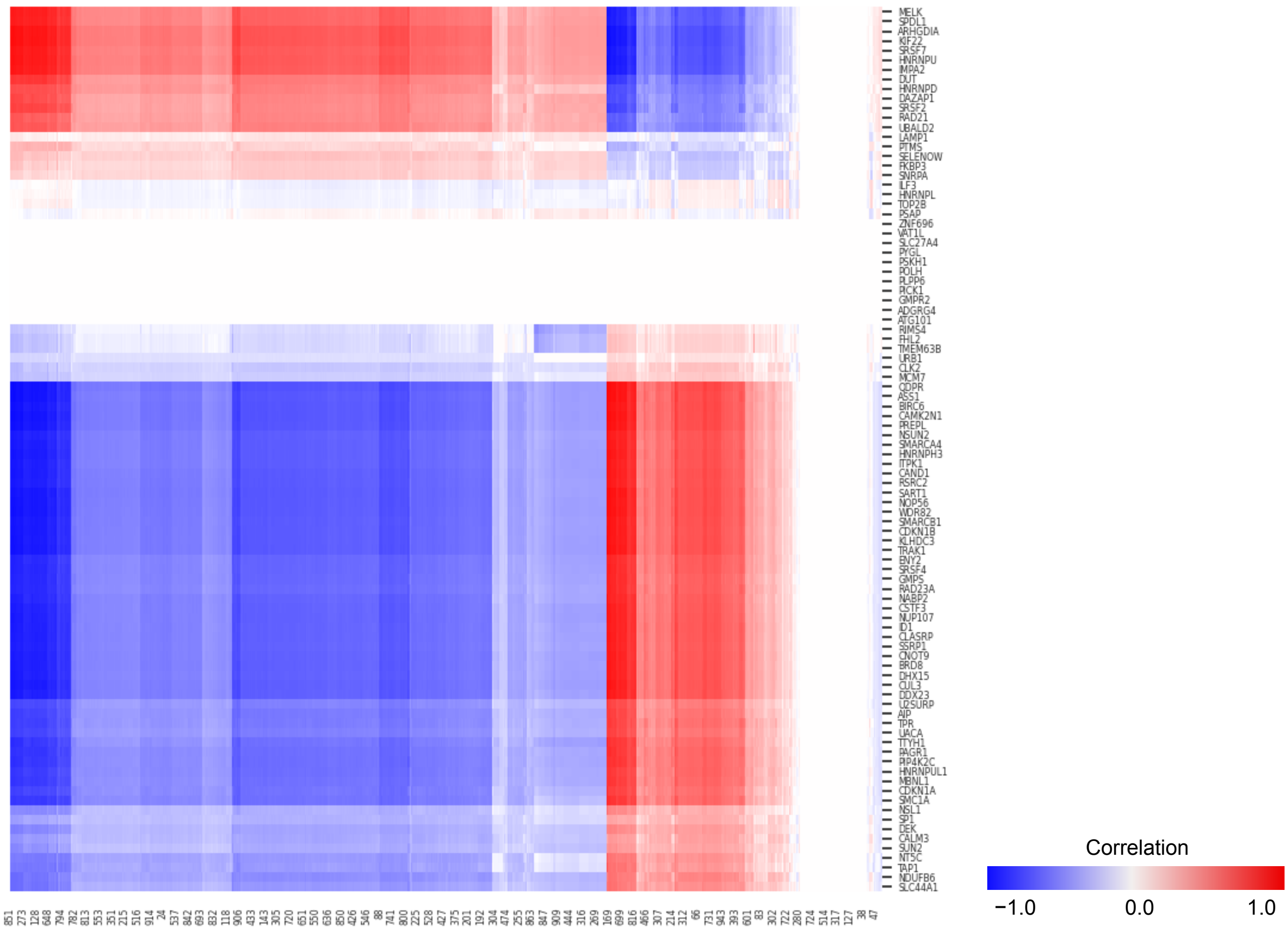

Correlation
−1.0
0.0
1.0
MELK
SPDL1
ARHGDIA
KIF22
SRSF7
HNRNPU
IMPA2
DUT
HNRNPD
DAZAP1
SRSF2
RAD21
UBALD2
LAMP1
PTMS
SELENOW
FKBP3
SNRPA
ILF3
HNRNPL
TOP2B
PSAP
ZNF696
VAT1L
SLC27A4
PYGL
PSKH1
POLH
PLPP6
PICK1
GMPR2
ADGRG4
ATG101
RIMS4
FHL2
TMEM63B
URB1
CLK2
MCM7
QDPR
ASS1
BIRC6
CAMK2N1
PREPL
NSUN2
SMARCA4
HNRNPH3
ITPK1
CAND1
RSRC2
SART1
NOP56
WDR82
SMARCB1
CDKN1B
KLHDC3
TRAK1
ENY2
SRSF4
GMPS
RAD23A
NABP2
CSTF3
NUP107
ID1
CLASRP
SSRP1
CNOT9
BRD8
DHX15
CUL3
DDX23
U2SURP
AIP
TPR
UACA
TTYH1
PAGR1
PIP4K2C
HNRNPUL1
MBNL1
CDKN1A
SMC1A
NSL1
SP1
DEK
CALM3
SUN2
NT5C
TAP1
NDUFB6
SLC44A1

Supplementary Figure S10: Spearman’s correlation heatmap between the mean gating values from the spatial gating units in the Gene Expression Predictor Module of PENNE and the predicted expression levels of each high confidence gene across all patches. Random permutation of PCM images lead to no associations with organelle-specific gene programs.

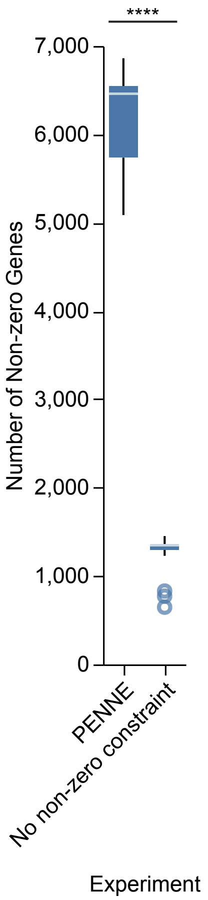

****
7,000
6,000
5,000
4,000
3,000
2,000
1,000
0
Number of Non-zero Genes
PENNE
No non-zero constraint
Experiment

Supplementary Figure S11: Box-and-whisker plot showing the number of non-zero predicted gene-expression levels across each sample for PENNE and the ablation model without the non-zero constraint loss. The number of non-zero predicted gene-expression levels is significantly higher for PENNE. Two-tailed Mann-Whitney $U$ test, ****: $p < 1.00 \times 10^{-4}$.

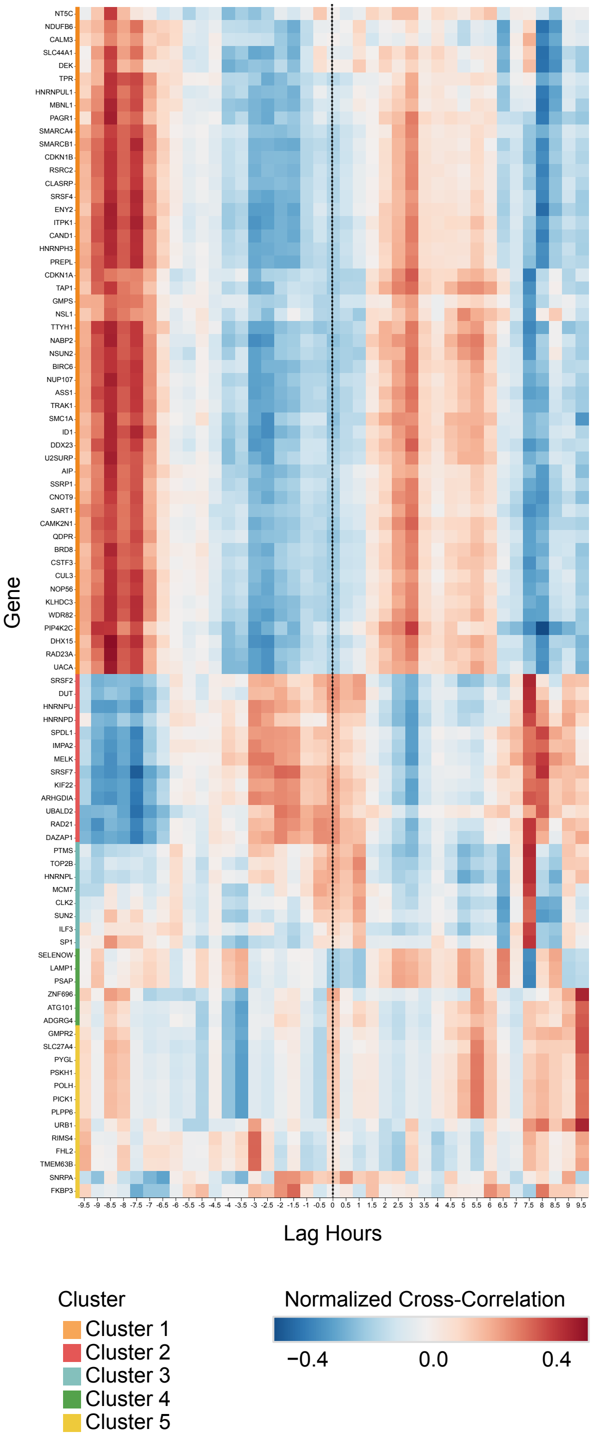
Gene
NT5C
NDUFB6
CALM3
SLC44A1
DEK
TPR
HNRNPUL1
MBNL1
PAGR1
SMARCA4
SMARCB1
CDKN1B
RSRC2
CLASRP
SRSF4
ENY2
ITPK1
CAND1
HNRNPH3
PREPL
CDKN1A
TAP1
GMPS
NSL1
TTYH1
NABP2
NSUN2
BIRC6
NUP107
ASS1
TRAK1
SMC1A
ID1
DDX23
U2SURP
AIP
SSRP1
CNOT9
SART1
CAMK2N1
QDPR
BRD8
CSTF3
CUL3
NOP56
KLHDC3
WDR82
PIP4K2C
DHX15
RAD23A
UACA
SRSF2
DUT
HNRNPU
HNRNPD
SPDL1
IMPA2
MELK
SRSF7
KIF22
ARHGDIA
UBALD2
RAD21
DAZAP1
PTMS
TOP2B
HNRNPL
MCM7
CLK2
SUN2
ILF3
SP1
SELENOW
LAMP1
PSAP
ZNF696
ATG101
ADGRG4
GMPR2
SLC27A4
PYGL
PSKH1
POLH
PICK1
PLPP6
URB1
RIMS4
FHL2
TMEM63B
SNRPA
FKBP3
-9.5 -9 -8.5 -8 -7.5 -7 -6.5 -6 -5.5 -5 -4.5 -4 -3.5 -3 -2.5 -2 -1.5 -1 -0.5 0 0.5 1 1.5 2 2.5 3 3.5 4 4.5 5 5.5 6 6.5 7 7.5 8 8.5 9 9.5
Lag Hours
Cluster
Cluster 1
Cluster 2
Cluster 3
Cluster 4
Cluster 5
Normalized Cross-Correlation
−0.4
0.0
0.4

Supplementary Figure S12: The complete cross-correlation heatmap showing the normalized cross-correlation between the predicted gene expression levels and geminin fluorescence levels across different time lags for each high-confidence gene.

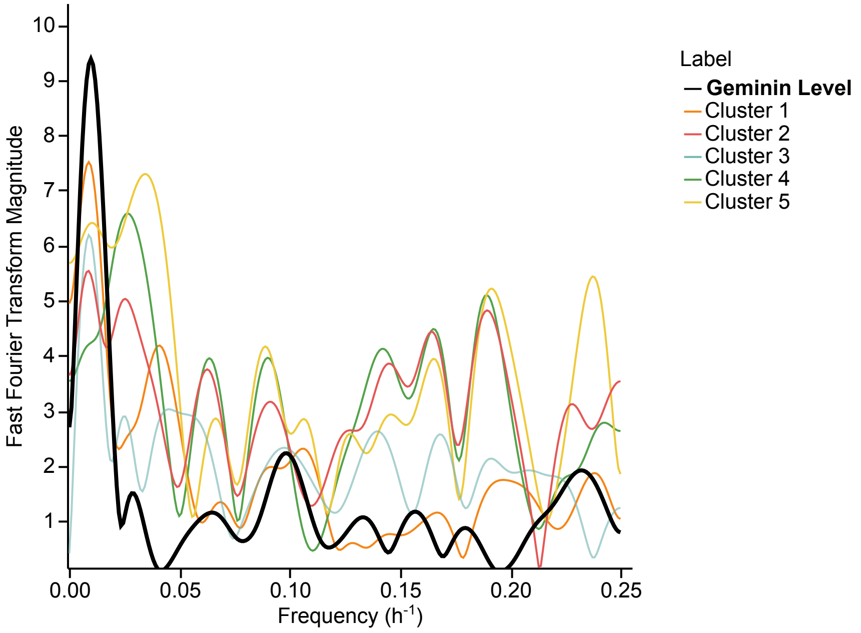

Fast Fourier Transform Magnitude
10
9
8
7
6
5
4
3
2
1
0.00
0.05
0.10
0.15
0.20
0.25
Frequency (h-1)
Label
Geminin Level
Cluster 1
Cluster 2
Cluster 3
Cluster 4
Cluster 5

Supplementary Figure S13: Fast Fourier transform analysis of the cross-correlation, identifying the dominant frequencies of the temporal correlation between the predicted gene-expression levels and the geminin fluorescence levels across different time lags for each cluster using MCF10A cells irradiated with 5Gy of radiation. The cell-cycle frequency decreases after irradiation, consistent with the expected cell-cycle arrest after radiation damage.

#### Supplementary Tables

Supplementary Table S1: Ablation study with different components of PENNE removed. All values are for gene sample Spearman's correlations (mean ± std). The best performance out of all experiments is **bolded**.

| Experiment | All | Non-Zero | High Confidence |
|---|---|---|---|
| No adversarial domain adaptation | **0.4760 ± 0.0204** | 0.4310 ± 0.0563 | 0.4300 ± 0.0495 |
| No SPAGHETTI | 0.41600 ± 0.00440 | 0.2140 ± 0.0237 | 0.202 ± 0.116 |
| No non-zero constraint loss | 0.4020 ± 0.0170 | 0.1230 ± 0.0180 | −0.0167 ± 0.0898 |
| No cell type and marker gene loss | 0.37200 ± 0.00510 | 0.23500 ± 0.00855 | 0.240 ± 0.166 |
| Complete PENNE | 0.4220 ± 0.0120 | **0.4570 ± 0.0672** | **0.5000 ± 0.0753** |
